# Metabolic imprinting drives epithelial memory during mucosal fungal infection

**DOI:** 10.1101/2025.07.11.664387

**Authors:** Jinendiran Sekar, Norma V. Solis, Jian Miao, Nicolas Millet, Bryce Tom, Derek Quintanilla, Aize Pellon, David L. Moyes, Joseph A. Gogos, Harry B. Rossiter, Scott G. Filler, Mihai G. Netea, Jennifer K. Yee, Marc Swidergall

## Abstract

Epithelial cells at barrier sites are emerging as active participants in innate immune memory, yet the underlying metabolic and epigenetic mechanisms remain unclear. Here, we uncover a previously unrecognized form of trained immunity in oral epithelial cells that enhances protection against fungal infection. Using a mouse model, we show that mucosal exposure to *Candida albicans* confers sustained protective memory that is independent of adaptive immunity and myeloid cells. Mechanistically, mucosal memory is driven by proline catabolism via proline dehydrogenase (Prodh) in epithelial cells, which sustains mitochondrial function, epigenetic remodeling, and promotes cytokine production upon secondary challenge. Unlike classical trained immunity in immune cells, epithelial memory is independent of glycolysis but partially sustained by fatty acid oxidation via carnitine palmitoyltransferase-I (CPT1). These findings uncover a distinct metabolic-epigenetic axis that underlines long-term epithelial memory in the oral mucosa and reveal novel non-hematopoietic mechanisms of mucosal defense against fungal pathogens.

## Introduction

Host defense relies on the finely tuned coordination of innate and adaptive immune responses to regulate commensal organisms and eliminate invading pathogens ^1,2^. While adaptive immunity is traditionally defined by its capacity to retain antigen-specific memory, accumulating evidence over the past decade demonstrates that innate immune cells can also develop memory-like features, a process termed trained immunity, which enables enhanced and rapid antigen-agnostic responses to secondary challenges with heterologous insults ^3–6^. Recent studies have extended this concept beyond immune cells, revealing that epithelial cells at barrier sites can also exhibit inflammatory memory, characterized by enhanced responsiveness following repeated injury or infection ^7–11^. Inflammatory memory is typically defined at the level of the chromatin landscape, where regions that gain accessibility during an initial inflammatory response remain open after resolution ^8,10^.

Metabolic reprogramming is a fundamental mechanism by which innate immune cells meet the energetic and biosynthetic demands of host defense. Trained immunity is sustained through long-lasting epigenetic modifications coupled with dynamic shifts in key metabolic pathways ^4^. For instance, trained immune cells undergo a metabolic shift from oxidative phosphorylation to glycolysis, which supports rapid ATP production and generates biosynthetic intermediates needed for cytokine production and epigenetic remodeling ^3,12^. Beyond glycolysis, fatty acid oxidation (FAO) contributes to innate immune memory through the provision of active lipid mediators, rather than ATP production ^13^. Additionally, amino acid metabolism plays a crucial role in immune function, serving both as an energy source and as a modulator of immune responses ^14^. In case of trained immunity induction, glutamine replenishment of the TCA cycle leads to accumulation of fumarate and subsequent inhibition of KDM5 histone demethylases ^15^. In addition, proline metabolism enhances resistance to bacterial infections ^16,17^. However, the regulatory role of amino acid metabolism, particularly proline catabolism in immune memory, remains unknown.

While distinct in their mechanisms, tissue-specific immunity and trained immunity are increasingly recognized as interconnected components of host defense. Recent advances, including the concept of integrated tissue immunity, propose that long-term protective responses arise from coordinated interactions among innate and adaptive immune cells, as well as tissue-resident non-hematopoietic cells ^18^. The oral mucosa represents a uniquely dynamic barrier that is constantly exposed to mechanical stress, food-borne antigens, and a diverse community of commensal microbes ^2,19^. Although the exact mechanisms of trained immunity induction in the oral mucosa remain unknown, imbalances in local immune responses are associated with increased susceptibility to disease ^20^.

*Candida albicans,* a central constituent of the human mycobiome, colonizes the oral mucosa in up to 75% of healthy individuals ^21^. Under conditions of local or systemic immunosuppression, however, *C. albicans* can transition to a pathogenic state and cause oral disease, termed oropharyngeal candidiasis (OPC) ^22^. In healthy individuals, *C. albicans* typically causes no harm, and its persistent colonization appears to be evolutionarily selected for its role in metabolic regulation and immune priming ^5,23–25^. During OPC, oral epithelial cells are continuously exposed to fungal components such as β-glucan, a key cell wall component of *C. albicans* known to induce trained immunity in immune cells ^3,5,6^. However, it remains unknown whether oral epithelial cells also retain memory of prior fungal exposure, such as through β-glucan recognition.

Here, we address critical gaps in understanding innate immune memory within the oral mucosa. We show that mucosal priming with *C. albicans* induces a protective memory in oral epithelial cells during reinfection, independent of adaptive immunity and innate immune cells. This memory depends on β-glucan–induced epigenetic remodeling and coordinated metabolic regulation. Specifically, it is fueled by proline catabolism via proline dehydrogenase (Prodh) and fatty acid oxidation through carnitine palmitoyltransferase 1 (CPT1). Inhibition of these pathways impairs cytokine responses and abrogates protection upon reinfection. These findings uncover a previously unrecognized, metabolically encoded epithelial memory program that is essential for mucosal antifungal defense.

## Results

### Mucosal priming promotes protective immunity during fungal reinfection

Tissue-specific immunity coordinates the interaction of resident immune cells, recruitment of their circulating counterparts and communication with neighboring non-immune epithelial, endothelial and stromal cells ^26,27^. Epithelial tissues at barrier sites establish inflammatory memory and enhanced responsiveness following repeated injury or infection ^7–10,28^. To explore how these mechanisms operate in oral mucosa tissues, we developed a mouse model of oral *C. albicans* reinfection (**Fig. 1A**). Wild-type mice pre-exposed to *C. albicans* exhibited significantly reduced fungal burden upon reinfection compared to naïve mice (**Fig. 1B**). Despite the enhanced protection during reinfection, cytokine and chemokine levels were similar betweensham- and infected mice before secondary infection (**Fig. S1**). However, primed mice showed enhanced inflammatory responses (**Fig. 1C; Fig. S2**) and increased neutrophil recruitment during early reinfection (**Fig. 1D**). To determine the contribution of adaptive immunity to enhanced protection during reinfection, we used *Rag1*^−/−^ mice, which lack functional T and B cells ^29^. Remarkably, primed *Rag1*^−/−^ mice still exhibited reduced oral fungal burden upon reinfection (**Fig. 1E**), indicating that enhanced protection is independent of adaptive immune mechanisms. β-glucan, a component of the fungal cell wall, has been shown to induce trained immunity in both animal models and human immune cells, resulting in enhanced immune responses to subsequent challenges ^30^. Therefore, we tested whether β-glucan exposure to the oral mucosa could mimic the protective effects of *C. albicans* primary infection (**Fig. S3A**). β-glucan primed mice exhibited enhanced fungal clearance upon *C. albicans* infection (**Fig. S3B**), even in the absence of adaptive immunity (**Fig. S3C**), mirroring the effects observed with live *C. albicans* priming. These data suggest that prior fungal exposure induces protective memory beyond adaptive immunity in the oral mucosa. Although trained immunity during invasive candidiasis is typically mediated by CCR2-monocytes and macrophages ^5,6^, our findings indicate that protection in the oral mucosa occurs independently of these cells. Mice lacking CCR2 exhibited enhanced resistance upon reinfection (**Fig. 1E**), suggesting that other host cell types contribute to mucosal immune memory. Given the enhanced neutrophil recruitment observed in primed mice during reinfection (**Fig. 1D**), we used antibody-mediated depletion in wild-type mice (**Fig. S4A**) to evaluate their role in mucosal protection. Neutrophils were required for fungal clearance during reinfection; however, their absence did not diminish the heightened inflammatory response (**Fig. 1F**; **Fig. S4-D**) suggesting that neutrophils act downstream of the amplified mucosal inflammatory environment during reinfection, responding to rather than initiating the enhanced immune response. Collectively, these results demonstrate that fungal-induced innate immune memory establishes an additional layer of protection within the oral mucosa, enhancing neutrophil-mediated pathogen clearance during secondary infection, independent of lymphocytes, monocytes and macrophages.

**Figure 1.**
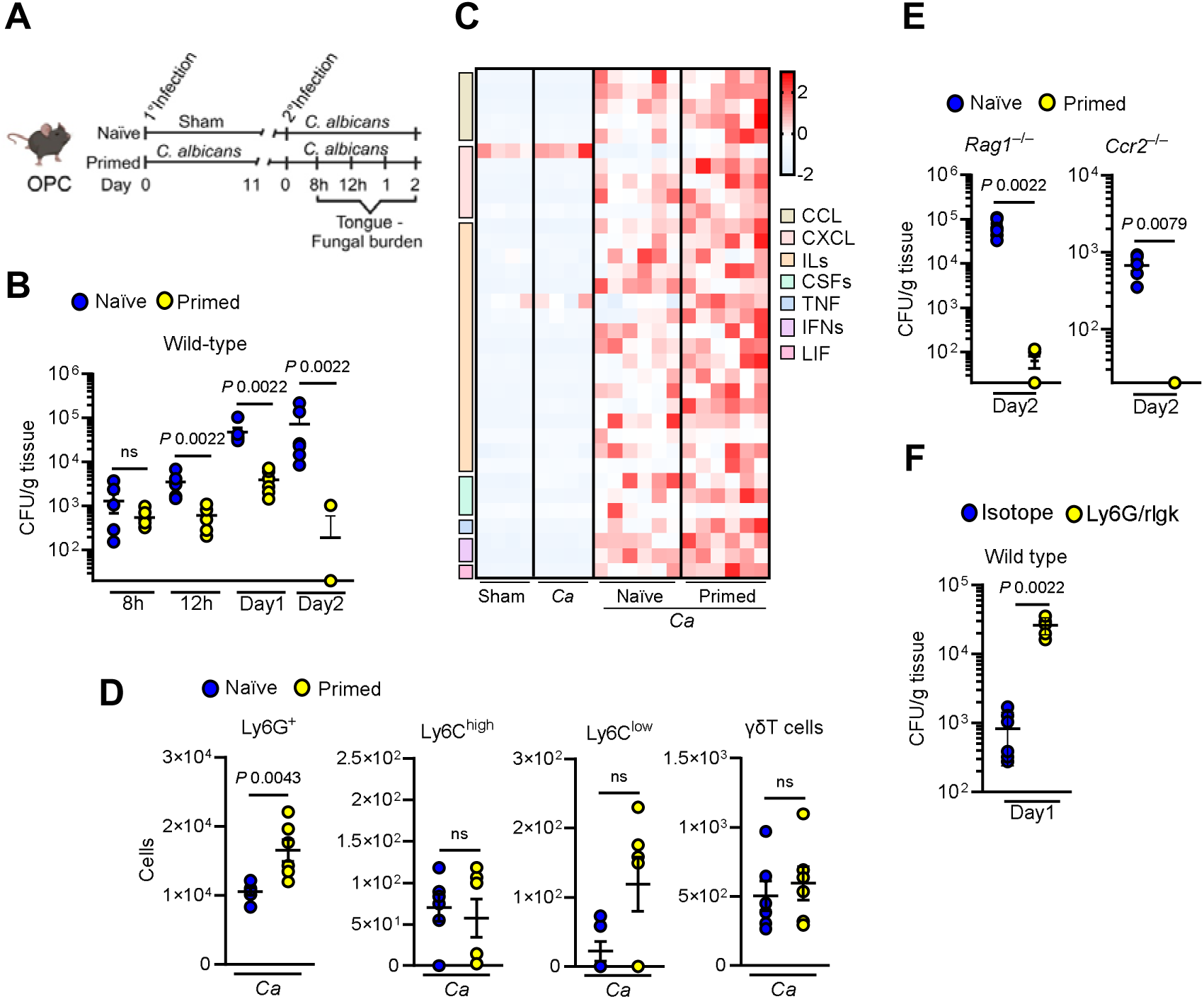
Mucosal priming enhances protection against oral reinfection. **A.** Schematic of the *C. albicans* reinfection model and experimental timeline for assessment. Created with BioRender.com **B**. Oral fungal burden in immunocompetent wild-type mice at indicated time points during reinfection. *N=6*; Two-tailed Mann–Whitney Test. The y-axis represents the limit of detection (20 colony-forming units [CFUs] per gram of tissue). **C.** Heat map of chemokine and cytokine levels in tongue homogenates from immunocompetent wild-type mice after 11 days of primary infection and 8h post-reinfection*. N=4-6*. **D.** Recruitment of immune cells to the tongue at 8h of reinfection. *N=6*; Two-tailed Mann–Whitney Test. **E.** Oral fungal burden in *Rag1*^−/−^ and *Ccr2*^−/−^ mice at 2 days post-reinfection. *N=6*; Two-tailed Mann–Whitney Test. **F.** Oral fungal burden in neutrophil-depleted mice 1 day post-reinfection. *N=6*; Two-tailed Mann–Whitney Test.

### β-glucan priming modulates inflammatory responses in epithelial cells

The oral epithelium serves as a critical barrier separating the host from the external environment and represents the first line of defense against fungal pathogens ^22^. As part of the innate mucosal antifungal response, epithelial cells release cytokines that recruit and activate neutrophils ^31^. Emerging evidence suggests that epithelial cells can acquire a form of inflammatory memory following prior stimulation ^8–11,32^. During oral fungal infection, epithelial cells are repeatedly exposed to the *C. albicans* cell wall component β-glucan ^33,34^. A key distinction between epithelial and immune cells lies in their response to this fungal ligand. While immune cells rapidly produce pro-inflammatory cytokines upon β-glucan recognition ^35,36^, epithelial cytokine levels remain unchanged following initial exposure ^37,38^. This observation suggests the existence of a ‘fail-safe’ mechanism in epithelial cells that limits unnecessary inflammation. Given the pivotal role of the oral epithelium in mucosal immunity, we hypothesized that enhanced responses upon *C. albicans* reinfection are mediated by β-glucan–induced memory in oral epithelial cells. To assess whether priming alters epithelial responsiveness, human oral epithelial cells were exposed with β-glucan or LPS, followed by cytokine measurement upon secondary challenge (**Fig. 2A**). Pre-exposure to β-glucan enhanced the production of specific pro-inflammatory cytokines following stimulation with *C. albicans* (**Fig. 2B**), even after an extended period (**Fig. S5A-B**). Additionally, epithelial cells primed with β-glucan showed an amplified cytokine response when subsequently challenged with LPS (**Fig. 2C**), suggesting enhanced response to heterologous stimuli. In contrast, LPS exposure followed by restimulation with LPS led to a tolerant response (**Fig. 2D**). In addition, priming with mannan, another component of the fungal cell wall ^39^, failed to enhance cytokine production upon subsequent stimulation with *C. albicans* (**Fig. S5C-D**). This suggests that the heightened immune response observed with β-glucan priming is not a general feature of all fungal microbe-associated molecular patterns, but rather a specific effect attributable to β-glucan. To determine the subsequent stimulus specificity, we tested cytokine production of epithelial cells following exposure to IL-17A alone or in combination with TNFα. Under these conditions, β-glucan priming did not increase CXCL8/IL-8 production (**Fig. S5E**), suggesting that the trained response is context and ligand-dependent. These findings support the concept that epithelial cells can retain and recall inflammatory memories from prior encounters. Given the functional parallels between trained oral epithelial and innate immune cells, we hypothesized that epigenetic remodeling may underlie β-glucan-induced epithelial memory.

**Figure 2.**
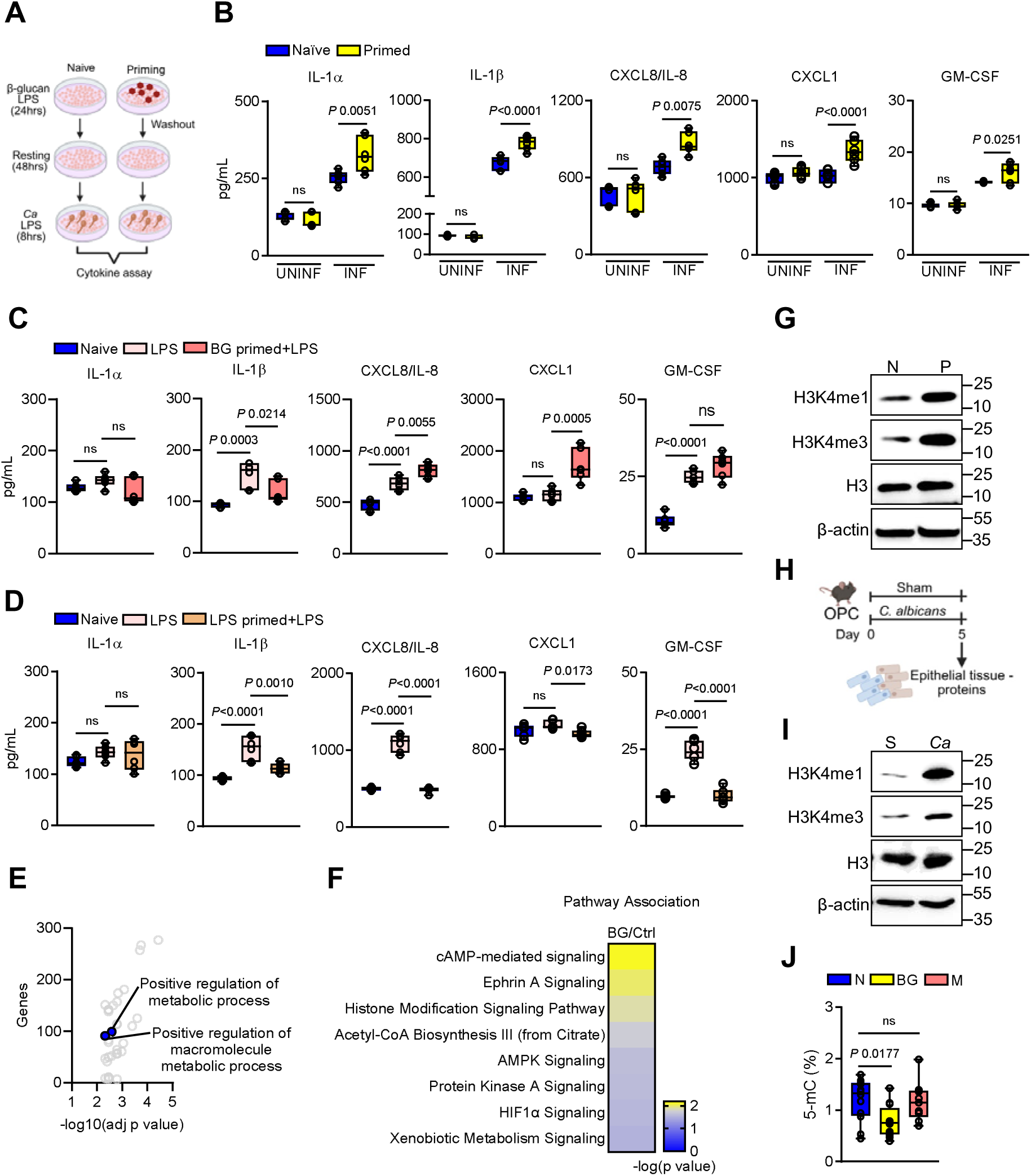
β-glucan priming induces epithelial memory. **A**. Schematic of the *in vitro* β-glucan priming model in human oral epithelial cells. Created with BioRender.com **B**. Chemokine and cytokine levels in supernatants from naïve and BG-primed epithelial cells after 8h of *C. albicans* infection*. N=6*; One-way ANOVA with Tukey’s multiple comparisons test. UNINF – uninfected, INF – Infected. **C**. Chemokine and cytokine levels in supernatants from naïve, LPS-treated, and BG-primed epithelial cells after LPS restimulation for 24h*. N=6*; One-way ANOVA with Tukey’s multiple comparisons test. **D**. Chemokine and cytokine levels in supernatants from naïve, LPS-treated, and LPS-primed epithelial cells after LPS restimulation for 24h. *N=6*; One-way ANOVA with Tukey’s multiple comparisons test. **E.** Volcano plot showing pathways associated with differentially accessible regions (DARs) identified by ATAC-seq after, analyzed using g: profiler. **F.** Stacked bar plot depicting canonical pathways significantly enriched between 24 h after β-glucan-stimulated and control oral epithelial cells, as determined by Ingenuity Pathway Analysis (IPA). *N=4*. **G.** Representative immunoblot showing histone methylation in β-glucan primed epithelial cells after 48h rest. N – naïve, P – β-glucan primed **H**. Schematic of fungal infection model and timeline of epithelial-enriched tissue collection. **I.** Representative immunoblot showing histone methylation in epithelial-enriched tissue from wildtype mice after primary infection. *N=3*. **J.** Global DNA methylation levels in epithelial cells primed with β-glucan and mannan for 24h followed by 48h resting. *N=6*; One-way ANOVA with Dunnett’s multiple comparisons test. N – naïve, BG – β-glucan, M – mannan.

To examine global chromatin accessibility, we performed an assay for transposase-accessible chromatin with sequencing (ATAC-seq) on epithelial cells during β-glucan stimulation (**Fig. S6A-C**). Gene Ontology (GO) analysis revealed that differentially accessible regions (DARs) were enriched in processes related to metabolic pathways (**Fig. 2E**). In parallel, Ingenuity Pathway Analysis (IPA) identified enrichment in pathways linked to histone modification, acetyl-CoA biosynthesis, and HIF-1α signaling (**Fig. 2F; Fig. S6D**). Immunoblotting validation of epigenetic changes revealed increased levels of histone methylation markers, including H3k4me1 and H3k4me3, in β-glucan-primed epithelial cells (**Fig. 2G; Fig. S7A**). Given that during acute OPC,

*C. albicans* is rapidly cleared from the oral cavity ^40,41^, epithelial-enriched tissues were collected to determine histone methylation upon primary infection (**Fig. 2H**). Similar to epithelial cells *in vitro*, epithelial-enriched tissues from mice exposed to *C. albicans* exhibit elevated levels of histone methylation (**Fig. 2I; Fig. S7B**). Moreover, global DNA methylation levels were reduced in β-glucan primed epithelial cells, a change not observed in cells exposed to mannan (**Fig. 2J**), further confirming the specificity of β-glucan-mediated epigenetic reprogramming. These findings demonstrate that β-glucan-induced long-lasting epigenetic modifications in epithelial cells likely contribute to their enhanced responsiveness during secondary challenge.

### β-glucan recognition induces proline catabolism in epithelial cells

In trained immune cells, enhanced responsiveness is often supported by altered metabolic regulation ^3,4,15,42^. Our chromatin accessibility profiling revealed enrichment of specific metabolic pathways in epithelial cells during β-glucan recognition (**Fig. 2E-F**), including pathways previously associated with trained immunity ^3,4,6^. To determine whether β-glucan recognition alters cellular metabolism, epithelial cells were stimulated at defined time points, and metabolite concentrations from different pathways were assessed by mass spectrometry (**Fig. 3A**). β-glucan recognition increased glucose and proline utilization, indicative of enhanced glycolytic activity and proline catabolism (**Fig. 3B**). Intracellular metabolic profiling further showed a shift in key pathways, including decreased proline and accumulation of glutamate, α-ketoglutarate, and citrate (**Fig. 3C-D**). These changes were corroborated by exposure of epithelial cells to heat-killed *C. albicans* yeast cells, which also induced glycolysis, proline catabolism, and accumulation of TCA cycle intermediates (**Fig. S8A-B**). To assess whether these metabolic changes are imprinted following β-glucan priming, epithelial cells were exposed to β-glucan for 24h and then allowed to return to homeostasis for 48h. Compared to naïve cells, primed epithelial cells exhibited persistent proline catabolism as evidenced by decreased proline levels and reduced glucose utilization (**Fig. S9A-B**), indicating metabolic imprinting that may facilitate rapid responses to secondary challenges. Additionally, β-glucan-primed cells exhibited increased mitochondrial oxidative phosphorylation (OXPHOS), including basal respiration, proton leak, ATP-linked respiration, and maximum respiration (**Fig. 3E-F; Fig. S10A-B**), suggesting enhanced mitochondrial respiration and oxidative phosphorylation. Consistent with *in vitro* observations, exposure of the oral mucosa in mice to *C. albicans* decreased proline and increased glutamate, α-ketoglutarate, and citrate levels (**Fig. 3G-H**). These changes persisted even after fungal clearance, suggesting that epithelial metabolic reprogramming may play a role in enhanced resistance upon reinfection.

**Figure 3.**
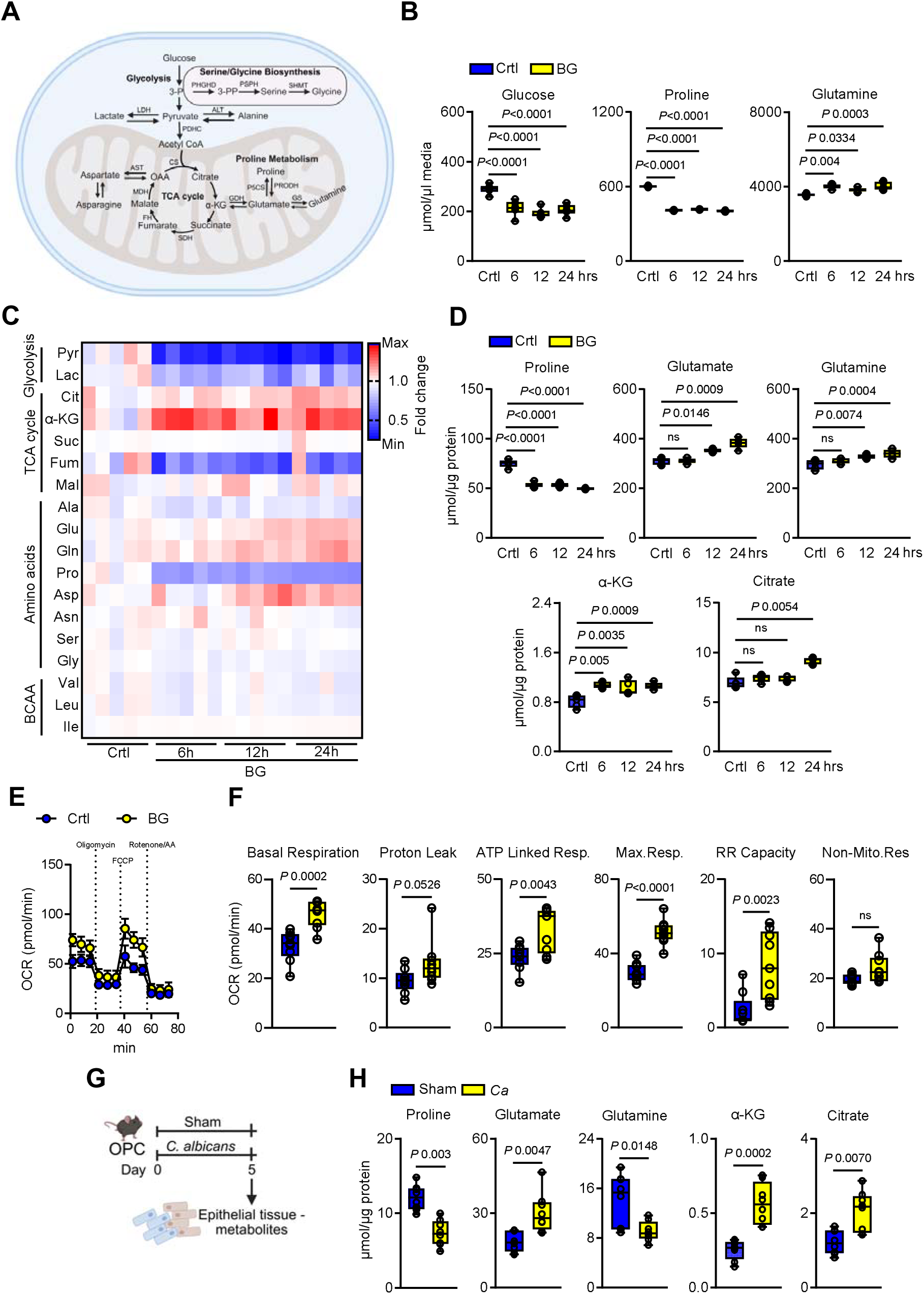
Recognition of β-glucan induces proline catabolism in epithelial cells. **A.** Schematic of key metabolites measured in epithelial cells involved in glycolysis, serine/glycine biosynthesis, the tricarboxylic acid (TCA) cycle, and proline metabolism following stimulation without or with β-glucan (BG). Created with BioRender.com **B.** Extracellular concentrations of glucose, proline, and glutamine in epithelial cells stimulated without or with β-glucan indicated time points. *N=6*; One-way ANOVA with Dunnett’s multiple comparisons test. **C**. Heat map showing the relative abundance of intracellular metabolites in epithelial cells stimulated without or with BG for the indicated time points. *N=6.* Abbreviations: Pyr – pyruvate, Lac – lactate; Cit – citrate, αKG – α-ketoglutarate, Suc – succinate, Fum – fumarate, Mal – malate, Ala – Alanine, Glu – glutamate, Gln – glutamine, Pro – proline, Asp – aspartate, Asn – asparagine, Ser – serine, Gly – glycine, Val – valine, Leu – leucine, IIe – isoleucine. **D**. Intracellular concentrations of selected metabolites in epithelial cells stimulated with BG for the indicated time points. *N=5.* One-way ANOVA with Dunnett’s multiple comparisons test. **E-F.** Mitochondrial oxidative function was quantified in epithelial cells stimulated without or with β–β-glucan for 24h, measured by Seahorse extracellular flux analysis. *N=9*; Unpaired student’s t-test. **G**. Schematic of the fungal infection and timeline of epithelial-enriched tissue metabolite measurement. Created with BioRender.com **H.** Concentrations of metabolites in epithelial-enriched tissues from immunocompetent wildtype mice after primary infection. *N=8.* Two-tailed Mann–Whitney Test.

### Proline catabolism via proline dehydrogenase promotes epithelial memory

β-glucan recognition and priming of epithelial cells induce a metabolic shift that favors proline catabolism over biosynthesis (**Fig. 3**). This reprogramming is evidenced by the upregulation of proline dehydrogenase (Prodh), the rate-limiting enzyme in proline catabolism, while the expression of pyrroline-5-carboxylate synthase (P5CS), which catalyzes proline biosynthesis, remains unchanged (**Fig. 4A-B; Fig. S11A-B**). Consistent with the *in vitro* data, *C. albicans* exposure increased Prodh expression in epithelial-enriched tissues without altering P5CS levels (**Fig. 4C; Fig. S11C**), supporting the concept that fungal recognition promotes proline catabolism. To determine whether proline catabolism is required for epithelial memory, we inhibited Prodh activity using tetrahydro-2-furoic acid (THFA) during β-glucan stimulation (**Fig. 4D**). Inhibition of Prodh resulted in increased proline levels but reduced downstream metabolites, including glutamate, α-ketoglutarate, and citrate (**Fig. 4E),** indicating that proline catabolism sustains TCA cycle intermediates. It was shown previously that proline catabolism fuels mitochondrial energy ^14,43^. Similarly, inhibition of Prodh reduced mitochondrial oxidative functions (**Fig. S12A-B**). Moreover, Prodh inhibition during β-glucan priming reduced the distinct cytokine responses to *C. albicans* infection (**Fig. 4F**). Proline metabolism regulates the epigenetic landscape of epithelial cells, including DNA methylation ^14,44–46^. Inhibition of Prodh increased the global DNA methylation in β-glucan primed epithelial cells (**Fig. 4G**), suggesting that proline catabolism is required for epigenetic rewiring in epithelial cells. Consistent with our *in vitro* findings, *Prodh*^−/−^ mice exhibited elevated proline levels and decreased TCA cycle intermediates in epithelial-enriched tissues post *C. albicans* infection (**Fig. 4H**). Importantly, *Prodh*^−/−^ mice maintained effective fungal control during primary infection (**Fig. S13A**). However, mice lacking Prodh exhibited increased susceptibility to reinfection and showed diminished inflammatory responses compared to their wild-type littermates. (**Fig. 4I-J; Fig. S13B-D**). Additionally, Prodh deficiency led to elevated global DNA methylation in epithelial-enriched tissue (**Fig. 4K**). Collectively, these data show that β-glucan-induced proline catabolism is a critical determinant of epithelial memory and antifungal immunity during reinfection.

**Figure 4.**
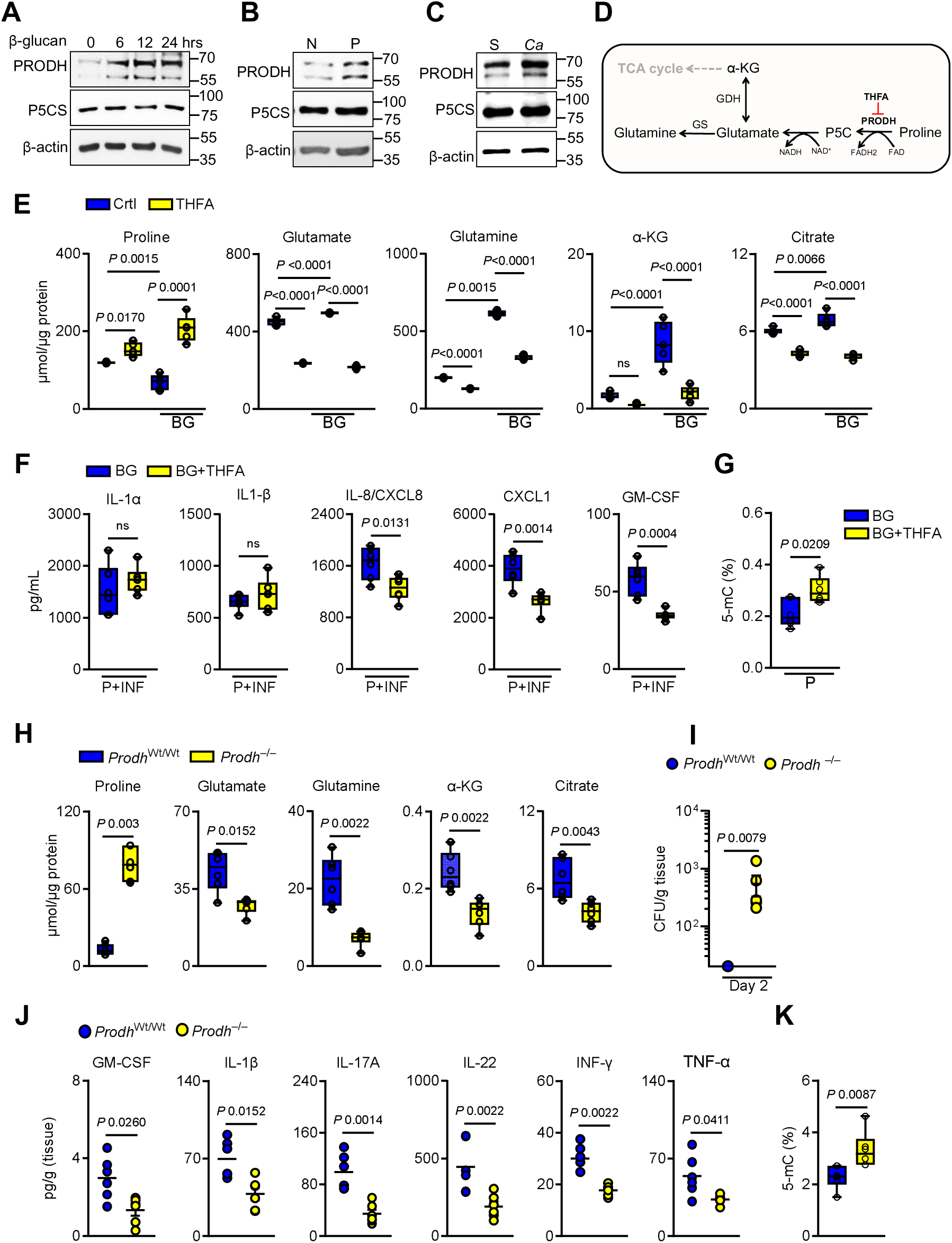
Epithelial memory is promoted by proline catabolism. **A-C.** Representative immunoblot analysis of proline dehydrogenase (PRODH; proline catabolism) and pyrroline-5-carboxylate synthase (P5CS; proline biosynthesis) expression in epithelial cells stimulated without or with β-glucan for the indicated times (**A**), in β-glucan primed epithelial cells after 48h resting (**B**), and in epithelial-enriched tissue from immunocompetent wildtype mice after 5 days of primary infection (**C**). N – naïve, P – β-glucan primed, S – Sham, *Ca* – *Candida albicans* **D.** Schematic diagram of the mitochondrial proline catabolism pathway. **E.** Intracellular metabolite concentrations in epithelial cells treated without or with 10mM tetrahydrofurfuryl acid (THFA; a proline dehydrogenase inhibitor) and stimulated with BG for 24h. *N=5.* One-way ANOVA with Tukey’s multiple comparisons test. **F.** Levels of chemokines and cytokines in culture supernatants of epithelial cells primed without or with THFA and BG after 8h of infection with *C. albicans. N=6.* Unpaired Student t-test. **G.** Global DNA methylation levels in epithelial cells trained with β-glucan and THFA for 24h, followed by 48h of resting in culture media. *N=6.* Two-tailed Mann–Whitney Test. **H.** Concentrations of metabolites in epithelial-enriched tissues from *Prodh*^wt/wt^ and *Prodh^−/−^*mice 5 days post-infection. **I.** Oral fungal burden in *Prodh*^wt/wt^ and *Prodh^−/−^* mice during reinfection. *N=6.* One-way ANOVA with Dunnett’s multiple comparisons test. The y-axis represents the limit of detection (20 CFUs/g of tissue). **J.** Proinflammatory cytokine response in tongue homogenates in *Prodh*^wt/wt^ and *Prodh^−/−^* mice upon reinfection. *N=6*; Two-tailed Mann–Whitney Test. **K.** Global DNA methylation levels in epithelial-enriched tissues from *Prodh*^wt/wt^ and *Prodh^−/−^* mice 5 days after primary infection.

### Protective epithelial memory is independent of glycolysis

β-glucan recognition in epithelial cells activates proline catabolism and glycolysis, highlighting coordinated metabolic remodeling during epithelial priming. Surprisingly, inhibition of Prodh during β-glucan exposure led to extracellular glucose accumulation (**Fig. 5A**), accompanied by reduced concentrations of glycolytic intermediates, including pyruvate and lactate (**Fig. 5B**), suggesting impaired glucose utilization and a potential metabolic reliance on proline catabolism to sustain glycolytic flux. Non-lethal levels of reactive oxygen species (ROS), byproducts of proline catabolism, can accumulate and stabilize hypoxia-inducible factor-1α (HIF-1α), a transcription factor that promotes glycolysis ^14,47,48^. Trained immune cells upregulate glycolysis via the mTOR-HIF-1α pathway ^3,4^. Therefore, we investigated whether proline catabolism regulates glucose metabolism through the ROS-HIF-1α axis. Inhibition of Prodh markedly reduced β-glucan-induced ROS accumulation, whereas β-glucan exposure alone increased ROS levels within 2h (**Fig. 5C**). Importantly, β-glucan recognition was not associated with cell death, as measured by lactate dehydrogenase (LDH) release (**Fig. S14A**). Consistently, Prodh inhibition also led to decreased expression of HIF-1α and glucose transporters Glut1 and Glut3 (**Fig. 5D; Fig. S14B**), supporting the role of proline catabolism as a driver of glycolysis (**Fig. 5E**). To assess the functional relevance of glycolysis in epithelial memory, we inhibited glycolysis using 2-deoxyglucose (2-DG) during β-glucan priming (**Fig. 5F**). 2-DG treatment suppressed pyruvate and lactate levels (**Fig. 5G**), while further depleting proline and TCA cycle intermediates (**Fig. 5H),** indicating metabolic compensation via accelerated proline catabolism. Functionally, inhibition of either HIF-1α or glycolysis enhanced the epithelial cytokine production in response to *C. albicans* (**Fig. 5I; Fig. S14C**), while decreasing the global DNA methylation (**Fig. 5J**). These findings suggest that protective epithelial memory persists independent of glycolysis, with proline catabolism acts as a central driver of both immune and epigenetic remodeling.

**Figure 5.**
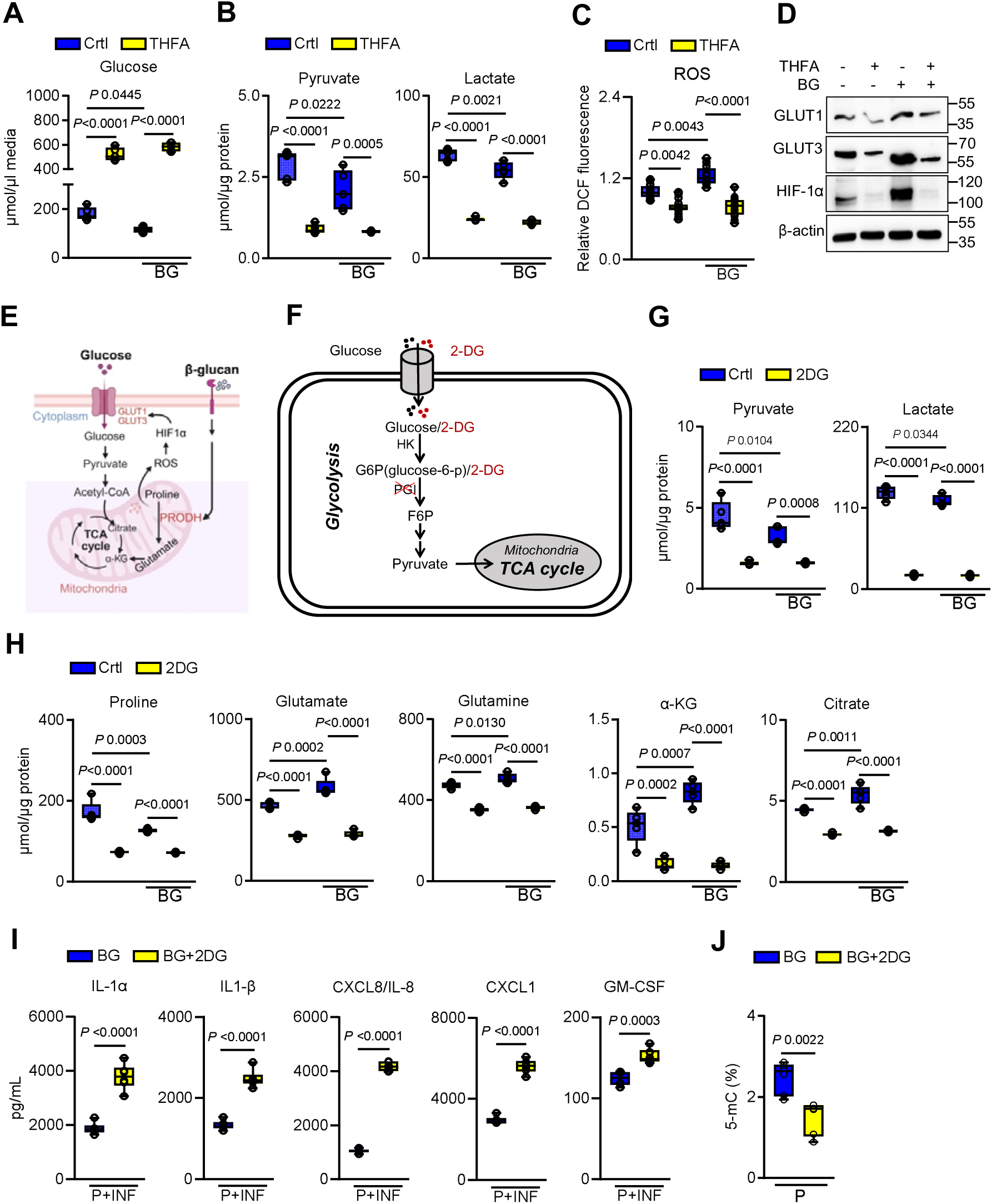
Protective epithelial memory occurs independently of glycolysis. **A.** Glucose concentrations in the media of epithelial cells treated without or with β-glucan (BG) and 10mM tetrahydrofurfuryl acid (THFA) for 24h. *N=5*. One-way ANOVA with Tukey’s multiple comparisons test. **B.** Intracellular pyruvate and lactate concentrations in epithelial cells treated without or with β-glucan and 10mM THFA for 24h. *N=5*. One-way ANOVA with Tukey’s multiple comparisons test. **C.** Intracellular reactive oxygen species levels in epithelial cells treated without or with THFA and BG for 2h. *N=9*; One-way ANOVA with Tukey’s multiple comparisons test. **D.** Representative immunoblot analysis of hypoxia-inducible factor 1-alpha (HIF-1α), glucose transporter 1-3 (GLUT1 and GLUT3) expression in epithelial cells treated without or with β-glucan and 10mM THFA for 24h. *N=3.* One-way ANOVA with Dunnett’s multiple comparisons test. **E.** Proposed schematic diagram of proline catabolism-mediated activation of glycolysis. Created with BioRender.com **F.** Schematic diagram of glycolysis and metabolite measurement approach. **G.** Glucose concentrations in media of epithelial cells treated without or with BG and 10mM 2-Deoxy-D-glucose (2DG; glycolysis inhibitor) for 24h. *N=5*. One-way ANOVA with Tukey’s multiple comparisons test. **H.** Intracellular metabolite concentrations in epithelial cells treated without or with BG and 10mM 2DG for 24h. *N=5*. One-way ANOVA with Tukey’s multiple comparisons test. **I.** Levels of chemokines and cytokines in culture supernatants of epithelial cells primed without or with 2DG/BG after 8h of infection with *C. albicans. N=6*. Unpaired Student t-test. **J.** Global DNA methylation levels in epithelial cells treated with β-glucan and 2DG for 24h followed by resting in culture media for 48h. *N=6*; Two-Tailed Mann–Whitney Test.

### Fatty acid oxidation partially contributes to epithelial memory

In immune cells, trained immunity is sustained by alterations of different metabolic pathways and epigenetic modifications ^49^. Among these, fatty acid oxidation (FAO) has emerged as a critical metabolic process regulating the induction and maintenance of trained immune responses ^4,13,15^. Proline catabolism and FAO are connected through their roles in cellular energy metabolism and redox homeostasis ^50^. Therefore, we determined fatty acid concentrations during β-glucan recognition and in primed epithelial cells. The levels of long-chain fatty acids (LCFAs) decreased, including palmitic, palmitoleic, stearic, oleic, and α-linolenic acids (**Fig. 6A-B**). Likewise, *C. albicans* exposure reduced LCFA levels in epithelial-enriched tissue after fungal clearance (**Fig. 6C**). Consistent with enhanced FAO, expression of carnitine palmitoyltransferase genes Cpt1 and Cpt2 were upregulated in β-glucan primed cells (**Fig. 6D-F; Fig. S15A-B**) and in epithelial-enriched tissue after *C. albicans* exposure (**Fig. 4G; Fig. S15C)**. Next, we inhibited CPT1 activity using etomoxir during β-glucan recognition. Targeted metabolome analysis revealed increased LCFA accumulation in etomoxir-treated cells (**Fig. 7A**). Functionally, CPT1 inhibition selectively reduced IL-1α and IL-1β production in primed epithelial cells upon *C. albicans* infection, while CXCL8/IL-8, CXCL1, and GM-CSF levels were elevated (**Fig. 7B**), indicating a partial dependence of the enhanced cytokine responses on FAO. FAO is known to regulate histone demethylases to enhance trained immunity in immune cells ^13,42,51^. Notably, inhibition of FAO decreased global DNA methylation in β-glucan-primed epithelial cells (**Fig.7C**) while reducing H3k4me1 and H3k4me3 levels (**Fig. 7D; Fig. S16A**). Consistent with increased distinct cytokines, inhibition of Prodh maintained histone methylation levels (**Fig. 16B-C**). To further link histone modifications to epithelial memory, we inhibited histone methyltransferases using 5′-methylthioadenosine (MTA) during β-glucan priming. MTA treatment attenuated IL-1α and IL-1β responses during *C. albicans* infection while increasing other cytokines (**Fig. 7E**). This suggests that FAO regulates histone methylation, while proline catabolism modifies DNA methylation to enhance distinct cytokines during secondary stimulation. Next, we determined whether both DNA and histone demethylation are required for epithelial memory by treating epithelial cells with 2-hydroxyglutarate (2-HG) during β-glucan training to inhibit α-ketoglutarate (α-KG)–dependent dioxygenases ^52,53^. 2-HG treatment suppressed cytokine responses to secondary *C. albicans* infection (**Figure 7F**), indicating that both DNA and histone demethylation are essential for the maintenance of enhanced responses during epithelial memory.

**Figure 6.**
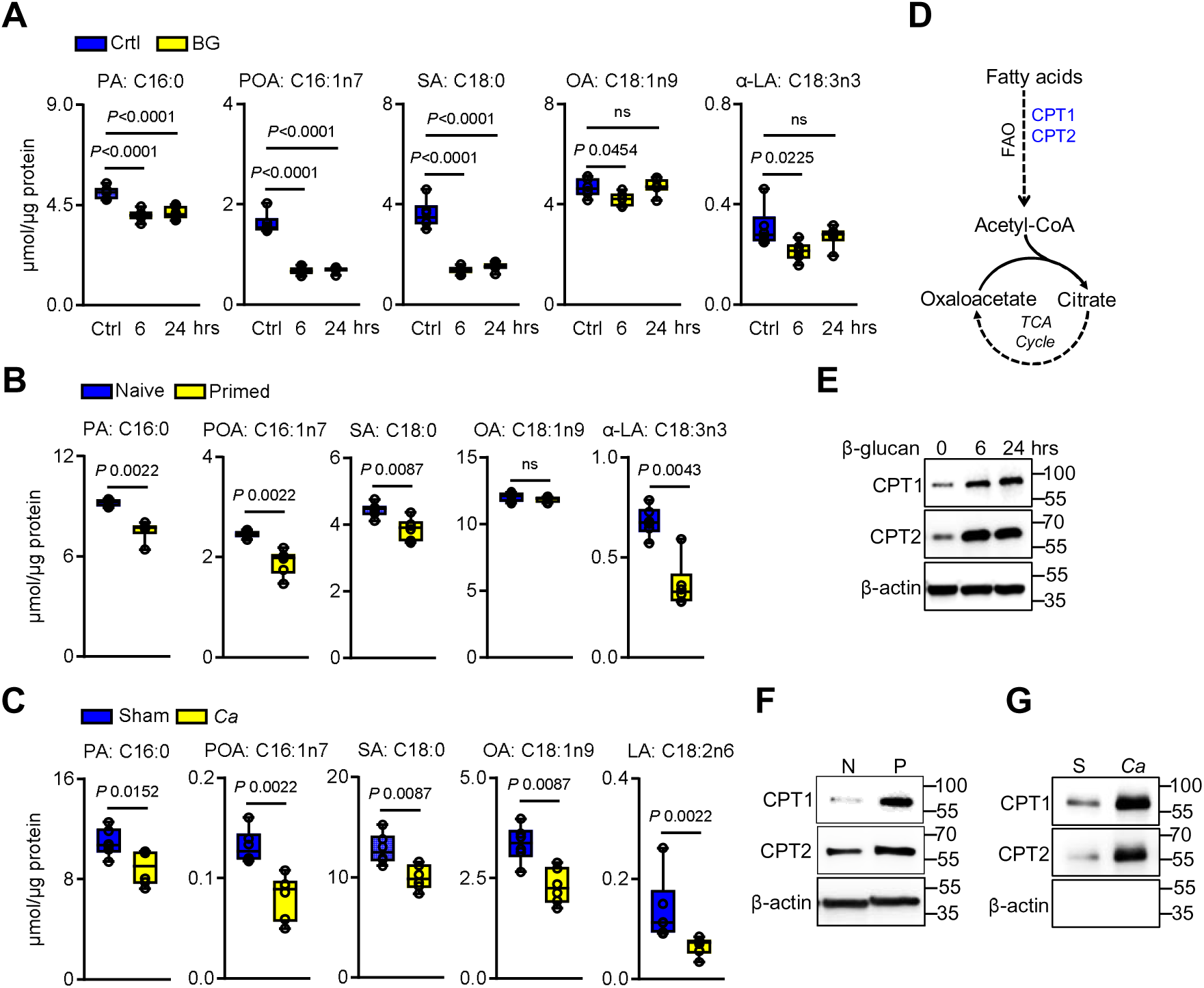
β-glucan recognition induces fatty acid oxidation in epithelial cells. **A.** Intracellular fatty acid concentrations in epithelial cells stimulated without or with β-glucan for 6 and 24h. *N=5*; One-way ANOVA with Dunnett’s multiple comparisons test. **B.** Intracellular fatty acid concentrations in BG-primed epithelial cells after 48h of rest. *N=5*; Two-tailed Mann– Whitney Test. **C.** Fatty acids concentrations in epithelial-enriched tissue from immunocompetent wild-type mice. *N=6.* Two-tailed Mann–Whitney Test. Abbreviations: PA:C16:0 – palmitic acid, POA:C16:1n7 – Palmitoleic acid, SA:C18:0 – Stearic acid, OA: C18:1n9 – oleic acid, α-LA:C18:3n3 – α-linoleic acid and LA:C18:2n6 – linoleic acid. **D.** Schematic of fatty acid oxidation and TCA cycle. **E-G.** Representative immunoblot analysis of carnitine palmitoyltransferase (CPT1/2) expression in epithelial cells stimulated without or with β-glucan for the indicated times (**E**), in β-glucan primed epithelial cells after 48h (**F**) and in epithelial-enriched tissue from immunocompetent wild-type mice after fungal clearance with primary infection(**G**).

**Figure 7.**
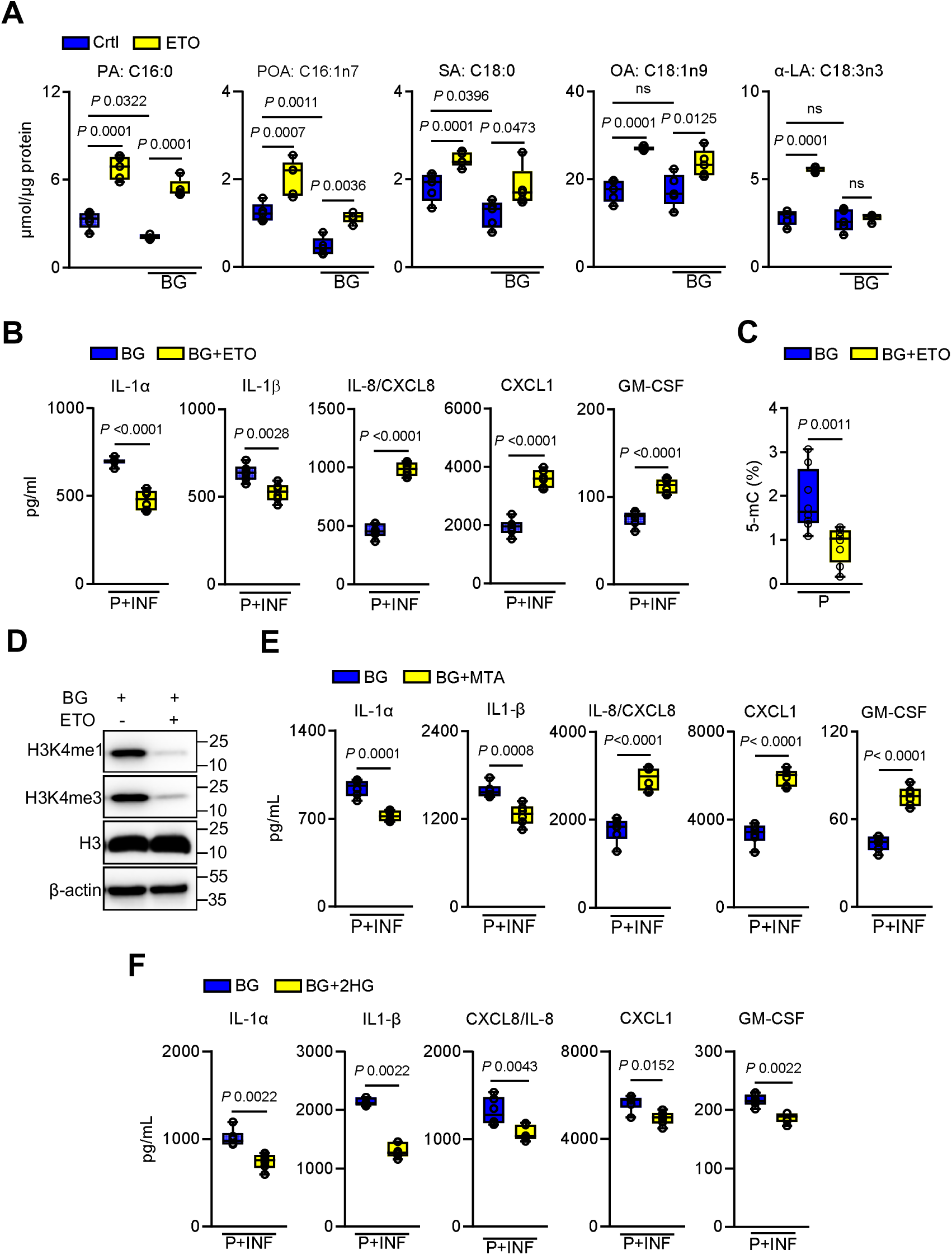
Epithelial memory is partially dependent on fatty acid oxidation. **A.** Fatty acid concentrations in epithelial cells treated without or with 100µM etomoxir (ETO; carnitine palmitoyltransferase inhibitor) and stimulated with BG for 24h. *N=5.* One-way ANOVA with Tukey’s multiple comparisons test. Abbreviations: PA:C16:0 – palmitic acid, POA:C16:1n7 – Palmitoleic acid, SA:C18:0 – Stearic acid, OA: C18:1n9 – oleic acid, and α-LA:C18:3n3 – α-linoleic acid. **B.** Levels of chemokines and cytokines in culture supernatants of epithelial cells primed without or with ETO/BG after 8h of infection with *C. albicans. N=6.* Unpaired student t-test. **C.** Global DNA methylation in epithelial cells treated with β-glucan and ETO for 24h and rested in culture media for 48h. *N=6.* Two-tailed Mann–Whitney Test. **D.** Representative immunoblot of histone methylation in primed epithelial cells without or with ETO/BG for 24h, followed by resting in culture media for 48h. **E.** Levels of chemokines and cytokines in culture supernatants of epithelial cells primed without or with the histone methyltrasferase inhibitor MTA (5-deoxy-5-methylthio-adenosine)/BG, followed by 8h of infection with *C. albicans. N=6.* Unpaired student’s t-test. **F.** Levels of chemokines and cytokines in the media supernatants of OECs primed without or with BG/10µM 2-hydroxyglutarate (2-HG) after 8h of infection with *C. albicans. N=6,* Un-paired t-test.

## Discussion

Epithelial barrier integrity is a primary determinant of resistance to mucosal fungal infections ^54,55^. Effective coordination between epithelial and immune cells maintains tissue homeostasis and promotes robust host defense ^34,56,57^. While trained immunity has traditionally been described in innate immune cells such as monocytes, macrophages, and NK cells ^58^, recent evidence suggests that non-hematopoietic cells, including epithelial stem and progenitor cells, can also develop memory-like responses following inflammatory or microbial exposure ^8,10,59^. In this study, we identify a previously unrecognized form of epithelial memory in the oral mucosa, with proline catabolism serving as a key driver of both immune and epigenetic remodeling, to enhance immune responses and consequently effector cell recruitment upon reinfection.

Recent studies demonstrate that epithelial stem cells retain inflammatory memory through epigenetic remodeling, enabling accelerated repair and enhanced immune responsiveness upon repeated insults ^8,10^. Similarly, airway epithelial cells acquire memory after bacterial exposure via chromatin remodeling ^7^. Consistent with this, we demonstrate that β-glucan-primed oral epithelial cells acquire a protective memory state accompanied by enhanced immune responsiveness to secondary insults. Tissue-specific metabolic reprogramming is recognized as a key modulator of host defense during infection ^9,11,60–62^. Here, we identified proline catabolism, mediated by Prodh, as a key metabolic driver of epithelial immune memory. Enhanced proline utilization supports mitochondrial function and increases levels of TCA cycle intermediates. Genetic or pharmacological inhibition of Prodh impairs mitochondrial function, suppresses cytokine production during reinfection, and increases global DNA methylation. These findings emphasize the tight connection between metabolic rate, epigenetic remodeling, and immune memory in epithelial cells.

Glycolysis is a hallmark of metabolic activation in immune cells, fueling cytokine production and antimicrobial function during infection ^11,48^. In immune cells, trained immunity depends on enhanced glycolytic flux ^3,4^. Unexpectedly, our results reveal that glycolysis is dispensable for the establishment and maintenance of epithelial immune memory. Inhibition of Prodh attenuates ROS generation, HIF-1α stabilization, and glucose utilization, suggesting that proline oxidation sustains glycolytic signaling without engaging classical glycolysis ^63,64^. Remarkably, glycolysis inhibition further enhances proline oxidation and cytokine production, highlighting the metabolic flexibility of primed epithelial cells. This is reminiscent of adaptive metabolic regulation in immune cells, which shift to FAO and amino acid metabolism when glycolysis is constrained ^15^. Although keratinocytes rely on glycolysis ^11^, our data reveal a unique metabolic reliance in oral epithelial cells on proline catabolism, which compensates for a lack of glycolytic activity during immune memory. Our findings also establish epithelial metabolic reprogramming with epigenetic plasticity. Inhibition of glycolysis reduced global DNA methylation in primed epithelial cells, consistent with prior studies linking metabolism to DNA methylation and histone modifications^3,4,15^.In this context, proline catabolism supports the epigenetic adaptation underlying epithelial immune memory independently of glycolysis.

Fatty acid metabolism is a well-established contributor to trained immunity in immune cells ^4,13^, but its role in epithelial memory remains less defined. Our results reveal that β-glucan priming and *C. albicans* exposure induce FAO via enhancement of the CPT system. Pharmacological inhibition of CPT1 suppresses IL-1α/β production during reinfection and decreases histone methylation marks, H3K4me1/3, suggesting a partial role for FAO in epithelial memory. Our findings align with FAO-dependent memory via epigenetic modifications observed in myeloid cells, stem cells, and airway epithelium ^13,42,51,65^. Moreover, epithelial cells maintain the memory of prior infection with *Streptococcus pneumonia* via specific H3k4me2 histone modification ^9^. We further show that proline catabolism-derived glutamate contributes to α-KG accumulation, fueling the TCA cycle and promoting epithelial immune memory. α-KG serves as a cofactor for α-KG–dependent dioxygenases that mediate active DNA and histone demethylation ^4^. Inhibition of these enzymes with 2-HG reduced cytokine production in primed epithelial cells following *C. albicans* exposure. These findings emphasize that epithelial memory relies on a metabolic-epigenetic axis driven primarily by proline catabolism and partially supported by FAO.

In conclusion, our findings redefine the oral epithelium as an active participant in innate immune memory, capable of mounting enhanced responses to secondary insults through metabolic and epigenetic reprogramming. Unlike classical trained immunity in immune cells, which relies on glycolysis, epithelial memory is driven by proline catabolism and supported by FAO. This metabolically encoded memory program enables rapid cytokine production and effector cell recruitment upon reinfection, offering a previously unrecognized mechanism of mucosal defense. By uncovering a central role for amino acid metabolism in non-hematopoietic cells, our study expands the paradigm of trained immunity and highlights epithelial tissues as dynamic sites of immunological adaptation with broad relevance for recurrent infections, mucosal homeostasis, and therapeutic targeting of tissue-resident memory responses.

## Acknowledgments

We thank Amy H Attaway (Cleaveland Clinic) for guidance with the ATAC sequencing experiments, Salomé LeibundGut-Landmann (University of Zurich) for the *Candida albicans* strain CA101, James G. Rheinwald (Dana-Farber/Harvard Cancer Center) for providing the OKF6/TERT-2 cell line, and members of the Division of Infectious Diseases at Harbor-UCLA Medical Center for critical suggestions.

## Author Contributions

Conceptualization: JS, MS

Methodology and Analysis: JS, NVS, HR, AP, DLM, JAG, DQ, JKY, MS

Investigation: JS, NVS, JM, NM, BT, JKY, MS

Funding acquisition: MS

Project administration: MS

Supervision: MS

Writing – original draft: JS

Writing – review & editing: JS, NVS, JM, NM, BT, DQ, AP, DLM, JAG, HR, SGF, MGN, JKY, MS

## Data Availability

The authors declare that the data supporting the findings of this study are available within the paper and its Supplementary Information files. ATAC-seq data have been deposited in the NCBI BioProject database under accession numbers (Accession numbers will be provided upon acceptance of the manuscript for publication).

## METHODS

### EXPERIMENTAL MODEL AND SUBJECT DETAILS

#### Chemicals

All reagents were obtained from Sigma Aldrich (Missouri, USA), and all antibodies were purchased from Cell Signaling Technologies (Massachusetts, USA) and Proteintech (Illinois, USA) unless otherwise stated. Keratinocyte Serum-Free media (KSFM) with supplements, Dulbecco Modified Eagle’s Medium (DMEM), Ham’s F-12 medium, and LC-MS grade organic solvents (methanol and ethyl-acetate) were purchased from ThermoFisher Scientific (Massachusetts, USA). Stable isotope-labeled compounds were purchased from CDN Isotopes (Montreal, Canada) and Cambridge Isotopes Laboratories (Massachusetts, USA). The derivatization reagent MTBSTFA+1%TBDMS was purchased from Restek (Pennsylvania, USA).

#### Ethics statement

All animal procedures were approved by the Institutional Animal Care and Use Committee (IACUC) at the Lundquist Institute for Biomedical Innovation at Harbor-UCLA Medical Center. All experiments were conducted in accordance with IACUC ethical guidelines, ensuring the use of appropriate anesthetics, humane endpoints, and measures to minimize animal distress.

#### Fungal strains and cell lines

*Candida albicans* strains SC5314 and CA101 were used in the experiments and cultured as previously described. The OKF6/TERT-2 oral epithelial cell line was cultured in a keratinocyte serum-free medium as previously described ^33^. OKF6/TERT-2 cells were authenticated by RNA-Seq (Conti et al., 2016), and tested for mycoplasma contamination before use.

#### Mouse model of oropharyngeal candidiasis

For animal studies, age- and sex-matched mice were used. Animals were bred and housed under pathogen-free conditions at the Lundquist Institute for Biomedical Innovation at Harbor-UCLA Medical Center. Mice were randomly assigned to the different treatment groups. Experiments were performed using homozygous *Prodh*^−/−^ and their wild-type *Prodh*^wt/wt^ littermates. The generation of *Prodh*^−/−^ mice has been as described previously ^66^. C57BL/6J, *Rag1*^−/−^ (B6.129S7-*Rag1^tm1Mom^*/J; Strain# 002216; non-leaky) ^29^ and *Ccr2^−/−^* (*B6.129S4-Ccr2^tm1Ifc^*/J; Strain#:004999) ^67^ mice were purchased from The Jackson Laboratory. All mice were cohoused for at least 1 week before experiments. Oropharyngeal candidiasis (OPC) was induced in all mouse strains as previously described ^40,68^. The *C. albicans* SC5314 strain was serially passaged 3 times in YPD broth, grown at 30 °C at 200 rpm for 16–24 h at each passage ^69^. Cells were harvested by centrifugation, washed twice with PBS, and counted using a hemacytometer. For infection, 7–8 week-old male and female mice were sedated, and a swab saturated with 2 × 10^7^ *C. albicans* cells was placed sublingually for 75 min. Using the same procedure, mice were re-infected 11 days after the primary infection. For antibody depletion experiments we modoifed a protocol used by Boivin and collaugues ^70^, wild-type mice were treated intraperitoneally on day −1 and 0 with 50 μg of Anti-Ly6G (clone 1A8, #BP0075-1) and anti-rat Kappa Immunoglobulin Light Chain (clone MAR18.5, #BE0122) and corresponding isotype controls (##BE0089 and #BE0085). Mice were sacrificed at 8 h, 12 h, and 1- and 2-days post-infection. For colony-forming unit (CFU) enumeration, tongues were harvested, weighed, homogenized and quantitatively cultured. Researchers were not blinded to experimental groups, as the endpoints (fungal burden and cytokine levels) were objective and based on measurable disease severity.

#### ATAC-seq analysis

To assess genome-wide chromatin accessibility, an Assay for Transposase-Accessible Chromatin with high-throughput sequencing (ATAC-seq) was performed. Briefly, OKF6/TERT-2 cells were stimulated with depleted zymosan for 24h and harvested, cryopreserved in culture media containing FBS and 5% DMSO, and shipped to Admera Health for ATAC-seq processing. Upon arrival, frozen cells were thawed, washed with PBS, and subjected to tagmentation as previously described ^71^. Cells were resuspended in ATAC lysis buffer and tagmented using the Nextera Library Prep Kit (Illumina). Tagmented DNA was purified with the MiniElute Reaction Clean-up Kit (Qiagen, Hilden, Germany) and subsequently amplified with barcode primers. Library quality and quantity were assessed using the Qubit 2.0 DNA HS Assay (ThermoFisher, Massachusetts, USA), TapeStation High Sensitivity D1000 Assay (Agilent Technologies, California, USA), and QuantStudio5 System (Applied Biosystems, California, USA). Libraries were pooled equimolarly based on quality control (QC) metrics and sequenced on an Illumina NovaSeq X Plus system (Illumina, California, USA) using a 150 bp paired-end (PE) configuration, generating 60 million PE reads (30 million reads per direction) per sample. Sequencing data were processed using the ENCODE ATAC-seq pipeline (https://github.com/ENCODE-DCC/atac-seq-pipeline), which includes adapter trimming, read mapping to the hg38 reference genome, duplicate filtering, and quality control. Deduplicated and mitochondria-filtered BAM files were generated, and peak calling was performed with MACS2 software (v2.2.7). Differential accessibility analysis was conducted using the R package DiffBind (v3.8.4), with narrow peak files as input. Peak annotation was carried out using the org.Hs.eg.db annotation package and ChIPseeker (v1.34.1). Genes associated with peaks located within promoter regions were selected for Gene Ontology (GO) and KEGG pathway enrichment analysis. Chromatin accessibility was assessed based on detailed output tables generated by Active Motif’s proprietary analysis software, which provided metrics including peak locations, sample comparisons, and gene annotations. Differentially accessible (DA) genes were classified as “open” or “closed” based on the directionality of DA peaks across gene regions (i.e., genes were classified as “open” if all associated DA peaks showed increased accessibility). Gene pathway analyses were performed using Ingenuity Pathway Analysis (IPA; Qiagen, Inc.) and g:Profiler (https://biit.cs.ut.ee/gprofiler/gost). For IPA, ATAC-seq DARs and their statistical significance values were uploaded, and canonical pathway enrichments were identified. g:Profiler results were visualized using Manhattan plots, with pathways grouped by data sources, including GO (Gene Ontology), REACTOME, TRANSFAC, miRTarBase, CORUM, and HP (Human Phenotype Ontology). The x-axis represents pathway groupings, and the y-axis represents adjusted enrichment p-values on –log10 scale.

#### Quantification of metabolites from human oral epithelial cells

Metabolite levels were measured in OKF6/TERT-2 cells using previously described methods ^72–74^. Briefly, cells were stimulated with 25µg/ml of depleted zymosan (InvivoGen; referred to as β-glucan) in keratinocyte serum-free (KSF) medium for 6, 12, or 24h. Alternatively, epithelial cells were exposed to heat-killed *C. albicans* strains SC5314 and CA101 for 24h. Following stimulation, cells were washed twice with PBS, and metabolites were extracted by adding 80% methanol and centrifugation at 15000g for 10min. Supernatants were collected and dried under a vacuum (Eppendorf). Dried samples were resuspended and spiked with 1mM of each of the following internal standards: [U-^13^C_3_]pyruvate, [^13^C_3_]lactate, [U-^13^C_6_]citrate, [U-^13^C_5_]α-ketoglutarate, [U-^13^C_4_]succinate, [U-^13^C_4_]fumarate and [U-^13^C_4_]malate. To convert carbonyl compounds (aldehydes and ketones) into their oxime derivatives, the samples were treated with 100µL of 5M hydroxylamine hydrochloride and 70µL of potassium hydroxide for 1.5h at 37°C. The pH of the samples was then adjusted to pH 2.0 and saturated with sodium chloride. Metabolites were extracted twice with ethyl acetate, and samples were dried under a nitrogen stream. For quantification of free amino acids, samples were spiked with 100µM of the following amino acids standards: [D_3_]serine, [^13^C_2_]glycine, [D_4_]alanine, [D_8_]valine, [U-^13^C_6_]leucine, [U-^13^C_6_]isoleucine, [D_3_]proline, [U-^13^C_5_]glutamate, [U-^13^C_5_]glutamine, [D3]aspartate and [D_3_]asparagine. The samples were dried under vacuum at ambient temperature. All dried samples were derivatized with 70μl of MTBSTFA with 1% TBDMS for TCA intermediates at 70°C for 1.5h, or amino acids at 80°C for 4h. Samples were centrifuged and transferred to GC vials for analysis. Metabolite analysis was performed on an Agilent 7890A Gas Chromatography (GC) system, coupled with an Agilent 5975C Mass Selective Detector (MSD) equipped with electron impact (EI) ionization. The GC was operated in splitless mode with a constant helium flow of 1mL/min. A 1µL injection of tert-butyldimethylsilyl derivatized metabolites was introduced into an HP-1MS column. The GC temperature program was as follows: initial temperature of 80°C, held for 1min, then increased at 10°C/min to 320°C and held at 320°C for 5min. For EI, the temperature program was as follows: start at 80°C, increase by 5°C/min to 250°C, then increase by 50°C/min to 300°C, and hold at 300°C for 5min. Mass spectra were acquired in selected ion monitoring (SIM) mode, and metabolites were quantified by measuring the peak areas of specific ions at their respective m/z ratios, using Agilent ChemStation software. The ions selected for metabolite quantification were: pyruvate m/z 274(M0) and 275(M+1); lactate m/z 261(M0) and 264 (M+3); citrate m/z 459(M0) and 463(M+6); α-KG m/z 446(M0) and 450(M+4); succinate m/z 289(M0) and 293(M+4); fumarate m/z 287(M0) and 289(M+2); and malate m/z 419(M0) and 422(M+4); alanine m/z 260(M0) and 264(M+4); glycine m/z 246(M0) and 248(M+2); valine m/z 288(M0) and 296(M+8); leucine m/z 274(M0) and 284(M+10); isoleucine m/z 302(M0) and 308(M+6); proline m/z 286(M0) and 289(M+3); glutamate m/z 432(M0) and 437(M+5); glutamate m/z 431(M0) and 436(M+5); serine m/z 390(M0) and 393 (M+3); aspartate m/z 418(M0) and 421(M+3) and asparagine m/z 417(M0) and 420(M+3). Metabolites were quantified by integrating peak areas from a sample and were normalized against the corresponding internal standards and the protein concentrations of the samples.

#### Epithelial-enriched metabolites

To measure metabolite concentrations in epithelial-enriched tissues, mice were orally infected with *C. albicans* SC5314 as described above. To minimize contamination by fungal metabolites, epithelial-enriched tissues were harvested at day 5 post-infection. Briefly, tongues were excised and placed in a 60mm petri dish, and epithelial tissues were isolated by tongue scraping, as previously described ^41^. Collected epithelial tissue was homogenized in 80% methanol using Matrix C tubes (MPBio) containing a ¼” ceramic bead (MPBio). After extraction, samples were dried, derivatized, and analyzed by GC-MS following the same procedure described above for cultured epithelial cells.

#### Quantification of glucose from oral epithelial cells

Glucose concentrations were measured using the aldonitrile-pentaacetate derivatization protocol previously described ^75,76^. Briefly, OKF6/TERT-2 cells were stimulated without or with β-glucan for the indicated time points. Culture supernatants were collected and clarified by centrifugation at 15000g for 10min. Samples were spiked with 1mM [U-^13^C_6_]glucose and dried under vacuum (Eppendorf). For derivatization, dried samples were treated with 100µl of 0.2M hydroxylamine hydrochloride dissolved in pyridine and incubated at 90°C for 40min. Subsequently, 100µl of acetic anhydride was added, and the samples were incubated at 90°C for an additional 60min. After derivatization, samples were dried again, resuspended in 100µl ethyl acetate, and transferred to GC-MS vials for analysis. Aldonitrile pentapropionate derivatives of glucose were analyzed in selected ion monitoring (SIM) mode, targeting m/z 314 (C1–C4), 319 (C1–C4), 242 (C1–C4), 225 (C1–C6), and 231 (C1–C6). Glucose concentrations were quantified by integrating peak areas, normalized to the corresponding internal standard, and adjusted based on the sample volume and cell number.

#### Quantifications for fatty acids from human oral epithelial cells and epithelial-enriched tissue

OKF6/TERT-2 cells were stimulated without or with depleted zymosan for 6 and 24h and washed twice with PBS. Acute fungal infection and isolation of epithelial-enriched tissue were described above. Cells and epithelial-enriched tissue were saponified overnight in 200 proof ethanol and 30% KOH (1:1, wt/vol) to release all fatty acids, including those from phospholipids, triglycerides, and cholesteryl esters, and then they were extracted with petroleum ether using a previously described method^77,78^. Fatty acids were derivatized as methyl esters using 0.5N methanolic HCl (Thermo Fisher Scientific). Derivatized fatty acids were analyzed by GC/MS using an Agilent 5975C selective mass detector connected to a model 7890A gas chromatography, with a Bpx70 column (30-m length, 250-μm diameter, 0.25-μm film thickness) from SGE (Austin, TX)^79^. The GC settings were: helium flow rate, 1 ml/min and initial oven temperature, 150°C, which was programmed to increase at 3°C/min to a final temperature of 221°C. Mass spectra were acquired using electron impact ionization and scan ion monitoring mode. Fatty acids were quantified by integrating peak areas from each sample and normalized against the corresponding internal standard (heptadecanoic acid-d3 (C17d3) and cell numbers.

#### Cytokine and chemokine measurements *in vivo*

To determine the whole tongue cytokine and chemokine protein concentrations, mice were orally infected with *C. albicans* as described above. Mice were sacrificed at indicated time points after post-infection. The tongues of infected mice were harvested, weighed, and homogenized to obtain tissue supernatants as we described previously ^33,34^. A multiplex bead-based assay was used to measure the concentration of multiple cytokines and chemokines simultaneously following the manufacturer’s protocols (Invitrogen).

#### Cytokine and chemokine measurements *in vitro*

To determine cytokine and chemokine levels in the culture supernatants using previously described methods ^33^. OKF6/TERT-2 cells were cultured in KSF media and stimulated for 24h with 25µg/ml depleted zymosan and 10ng/ml lipopolysaccharide (Sigma). To identify the metabolic pathways responsible for enhanced cytokines productions, cells were treated with various inhibitors: PRODH inhibitor (10mM, Cat No.185396, sigma), glycolysis inhibitor (10mM, Cat No.S4701, SelleckChem), CPT1 inhibitor (100µm; Cat No. S8244; SelleckChem), histone methylation inhibitor (100µM; Cat No. E1024; SelleckChem), and HIF-1α inhibitor (10µM; IDF-11774; SelleckChem). Cells were washed twice with PBS, then re-incubated in fresh KSF media and allowed to rest for 48h. Cells were infected with *C. albicans* SC5314 at a multiplicity of infection (MOI) of 5 for 8h. Alternatively, cells were stimulated with a combination of 50ng/ml IL-17A and 0.5ng/ml TNFα (Peprotech) for 8h to simulate inflammatory conditions. After 8h of infection or stimulation, culture supernatants were collected, clarified by centrifugation and stored in aliquots at −80°C for later analysis. The levels of proinflammatory mediators in culture supernatants were quantified using the Luminex multiplex assay (R&D Systems).

#### Flow cytometry of infiltrating leukocytes

Immune cells in the tongues were characterized as described previously ^69^. Mice were infected with *C. albicans* and harvested tongues after 8h of reinfection as described above. Single-cell suspensions from tongues were prepared using previously described methods. Briefly, tongues were finely minced and digested using 1ml digestion buffer (4.8 mg/ml collagenase IV, 200 μg/ml DNase I) at 37°C for 45 min with shaking. Digested tissue was passed through a 70-μm filter strainer and washed. Cells were resuspended in RPMI 1640 medium, and they were overlaid on 40% Percoll followed by centrifugation at 350×*g* for 30 min at 37°C. Cells were washed and stained with a LIVE/DEAD Fixable Blue viability kit (Thermo Fisher) for 15 min, washed, and resuspended in FACS buffer. Cells were then incubated with rat anti-mouse CD16/32 (2.4G2; BD Biosciences) for 15 min (1:100) in FACS buffer at 4 °C to block Fc receptors. For staining of surface antigens, cells were incubated with fluorochrome-conjugated (FITC, PE, PE-Cy7, allophycocyanin (APC), APC-Cy7, and Brilliant Violet 650) antibodies against mouse CD4 (GK1.5, BioLegend), CD11b (M1/70, Invitrogen), CD45 (QA17A26, BioLegend), TCRγ/δ (GL3, BioLegend), Ly6C (HK1.4, BioLegend), and Ly6G (1A8, Biolegend). All antibodies used for flow cytometry were diluted 1:100 ratio. The stained cells were analyzed on a FACSymphony A5 system (BD Biosciences), and data were acquired using FACS Diva (BD Biosciences) and analyzed with FlowJo software. Only single cells were analyzed, and cell numbers were quantified using PE-conjugated fluorescent beads (Spherotech). Single cells were identified as singlet’s CD45, CD11b^+^, Ly6G^+^, Ly6C^high^, Ly6C^low^, and γδT cells (**Figure S17**).

#### Quantification of global DNA methylation

OKF6/TERT-2 cells (3.5×10^4^) were plated in 6-well plates and incubated overnight at 37°C to allow the cells to settle. Cells were stimulated without or with 25µg/ml of depleted zymosan and mannan (Sigma) in KSF media, as described above. After 48h of resting, cells were trypsinized, and genomic DNA was isolated using FitAmp™ Blood and Cultured Cell DNA Extraction Kit (EpigenTek, New York, USA) according to the manufacturer’s instructions. The concentration of genomic DNA was measured using a NanoDrop Spectrophotometer (ThermoScientific, USA). Global DNA methylation levels were assessed using the MethylFlash™ Global DNA Methylation (5-mC) ELISA Easy Kit (EpigenTek, New York, USA), following the manufacturer’s protocols.

#### Mitochondrial oxidative function in intact oral epithelial cells

Mitochondrial function in intact OKF6/TERT-2 cells was assessed using Seahorse XF HS Mini Analyzer^72,74,80^. Briefly, cells were cultured in KSF media and stimulated without or with 25µg/ml depleted zymosan for 24h and rested for culture media for 48h as described above. One day prior to the assay, 6×10^4^ cells were grown in an 8-well Seahorse assay plate. Mitochondrial stress tests were performed using the manufacturer’s protocol. To measure ATP synthesis linked to oxidative phosphorylation, cells were treated with oligomycin. Maximum respiratory capacity (uncoupled respiration) was determined by adding FCCP, a protonophore that uncoupler’s mitochondrial oxidative phosphorylation. Non-mitochondrial respiration (residual oxygen consumption) was quantified by inhibiting mitochondrial complexes using rotenone (complex I inhibitor) and antimycin A (complex III inhibitor). All experiments were performed with biological replicates (n=6 or 9), and results were expressed as mean oxygen consumption rate.

#### ROS measurement

The fluorescent probe 2’,7’-dichlorodihydrofluorescein diacetate (DCF-DA; Fisher Scientific) was used to measure intracellular accumulation of reactive oxygen species (ROS) ^81^. Briefly, OKF6/TERT-2 cells (4×10^4^) were plated in 96-well plates and incubated overnight to settle down. Cells were incubated in a KSF media containing 10µM CM-H2DCFAD (Invitrogen) for 45 min, and then the medium was removed and replaced with fresh medium. Cells were treated with β-glucan or/and THFA in KSF media and immediately placed in the IncuCyte SX5 Live Cell Analysis System (Sartorius, Göttingen, Germany). ROS production was observed and automatically analyzed from acquired three images per well, taken every 30 min for a consecutive 24h. Data was analyzed with the Incucyte live cell analysis software.

#### Immunoblots

OKF6/TERT-2 cells (2×10^5^) grown in 24-well plates were switched to KSF media without supplements for 1h. Cells were then stimulated with 25µg/ml depleted zymosanfor various time points. After treatments, cells were lysed with ice-cold lysis buffer containing protease and phosphatase inhibitors (Thermo Scientific). Protein samples were prepared and separated by SDS-polyacrylamide gel electrophoresis. Proteins were transferred to polyvinylidene fluoride membranes (Bio-Rad, USA) by electroblotting. Phosphorylation and total protein levels were detected by immunoblotting using specific antibodies: H3K4me1 (Cat#; 7110795; 1:2000 dilution), H3K4me3 (Cat#; 9751T; 1:2000 dilution), histone H3 (Cat#; 4499T), PRODH/POX (Cat#: 22980, 1:2000 dilution), ALDH18A1/P5CS (Cat#: 17719; 1:2000 dilution), HIF-1α (Cat#; 36169T; 1:2000 dilution), GLUT1 (Cat#; 81463; 1:2000 dilution), GLUT2 (Cat#; 20403; 1:2000 dilution), CPT1B (Cat#; 41803T; 1:2000 dilution), CPT2 (Cat#; 52552T; 1:2000 dilution), and β-actin (8H10D10, #3700). Each experiment was performed at least three times. Immunoblots were developed using chemiluminescence and imaged with a C400 digital imager (Azure Biosystems, CA, USA). Protein expression levels were quantified by measuring the band intensity using ImageJ software. The β-actin was used as a loading control to normalize protein expression.

#### Statistical analysis

Except for the ATAC-seq experiment, all statistical analyses were performed using GraphPad Prism version 8.0. For pairwise comparisons of variables, a two-tailed unpaired Student’s t-test and Mann-Whitney were used. For multiple comparisons, a one-way analysis of variance (ANOVA) was conducted, followed by Tukey’s or Dunnett’s post-host test as appropriate. A value of *P* < 0.05 was considered statistically significant. Data are presented as the mean ± standard error of the mean (s.e.m.), with error bars indicating the variability of the data.

