## Supplemental Material for "Metabolic imprinting drives epithelial memory during mucosal fungal infection"

Supplementary Information Figure S1-S17

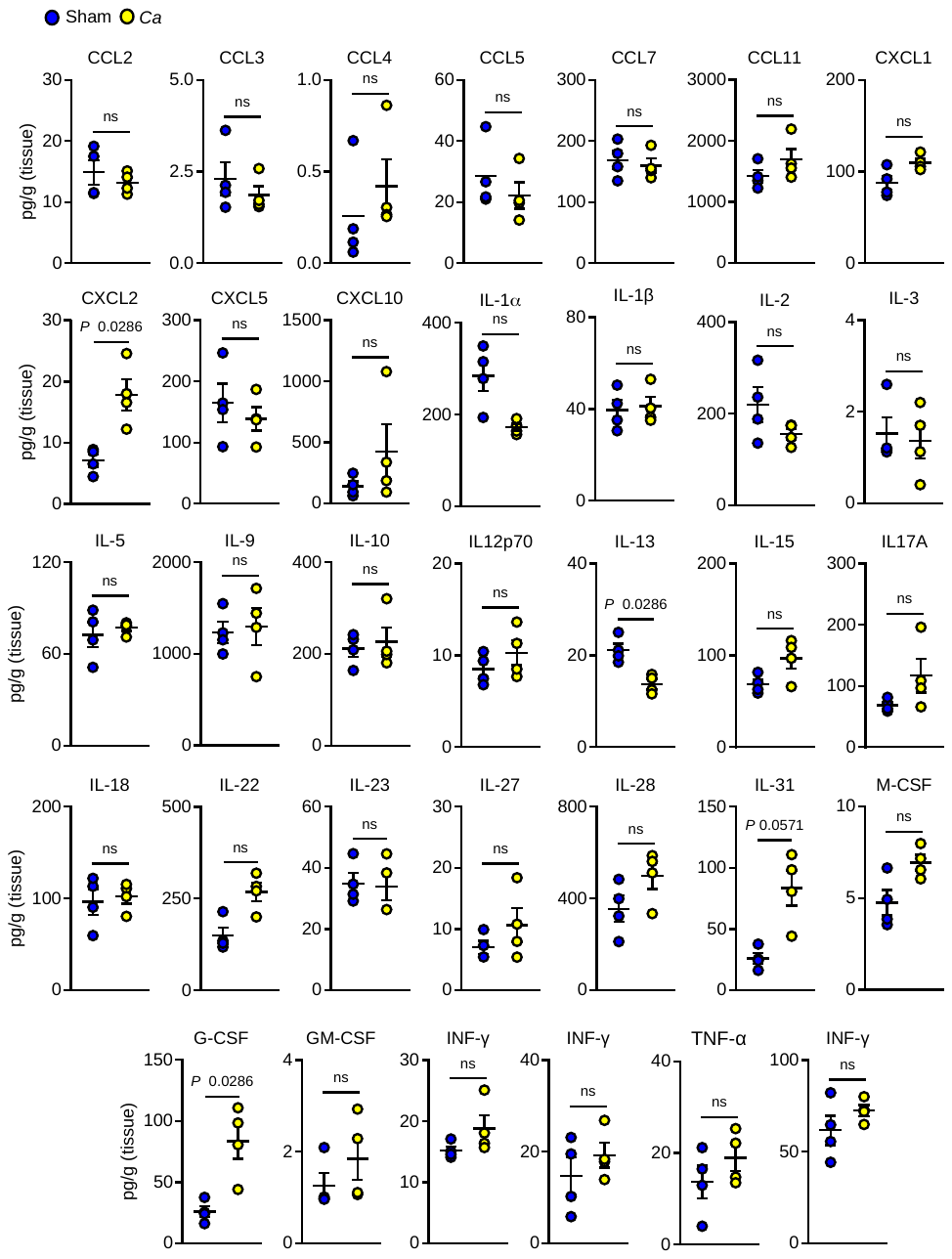

**Figure S1. Immune response in wild-type mice after 11 days of primary infection**. (Related to Figure 1). Cytokine and chemokine levels in tongue homogenates from immunocompetent wild-type mice were analyzed on day 11 after primary infection. *N=4*; Two-tailed Mann–Whitney Test.

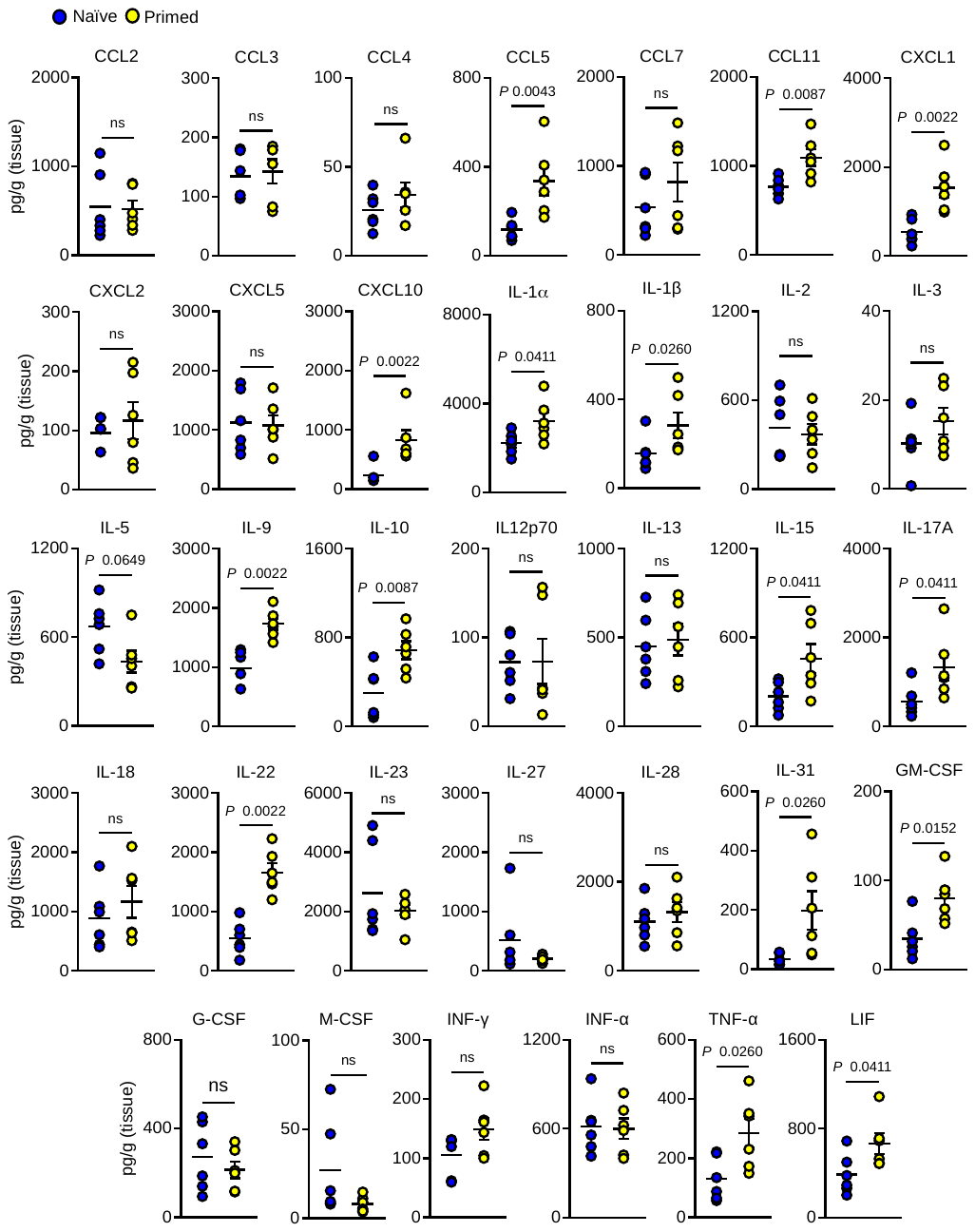

**Figure S2. Mucosal priming enhances proinflammatory cytokine response during reinfection.** (Related to Figure 1)**.** Levels of cytokines and chemokines in tongue homogenates from immunocompetent wild-type mice after 8h of post-reinfection. *N=6*. Two-tailed Mann–Whitney Test.

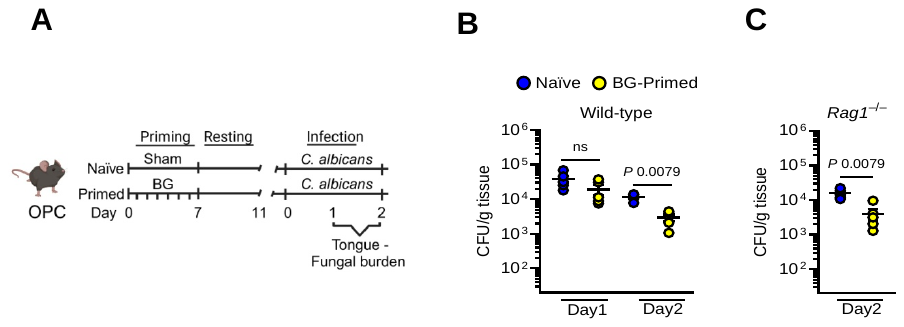

**Figure S3. Mucosal priming of β-glucan enhances protection during infection. A**. Schematic of β-glucan priming and experimental timeline for fungal burden assessment during infection. Created with BioRender.com **B**. Oral fungal burden in β-glucan-primed immunocompetent wild-type mice at 1 and 2 days post-reinfection. *N=6*; Two-tailed Mann–Whitney Test. **C.** Oral fungal burden in β-glucan-primed *Rag1*^–/–^ mice on day 2 postinfection. *N=6*; Two-tailed Mann–Whitney Test. The y-axis represents the limit of detection (20 CFU/g tissue).

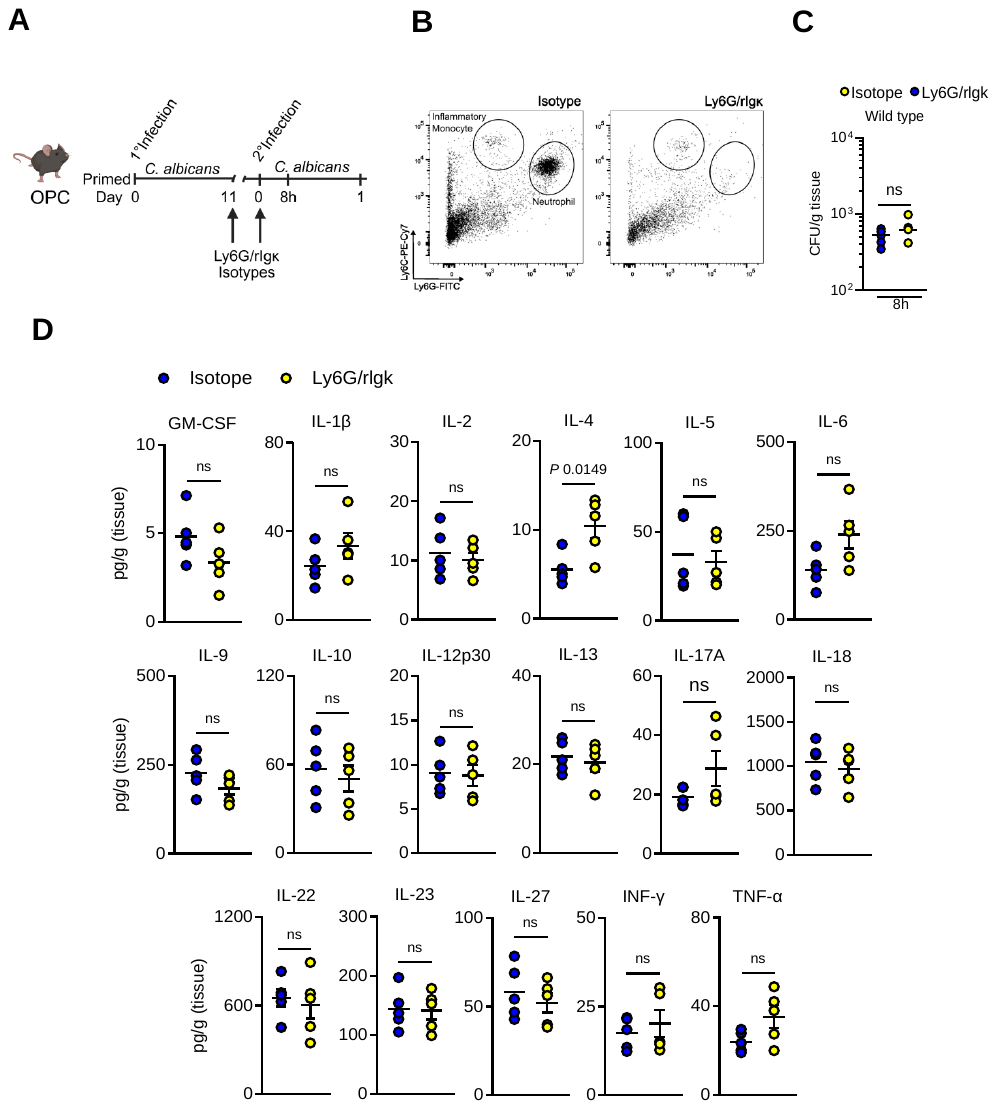

**Figure S4. Cytokine response unchanged in neutrophil-depleted mice during reinfection** (Related to Figure 1). **A.** Schematic illustrating neutrophil depletion strategy. Created with BioRender.com **B.** Gating strategy for neutrophils and inflammatory monocytes in the blood after 1 day of post-reinfection. **C.** Oral fungal burden in neutrophil-depleted mice after 8h of reinfection. *N=5*; Two-tailed Mann–Whitney Test. **D.** Cytokine and chemokine levels in tongue homogenates from neutrophil-depleted mice 8h after reinfection. *N=5*. Two-tailed Mann–Whitney Test.

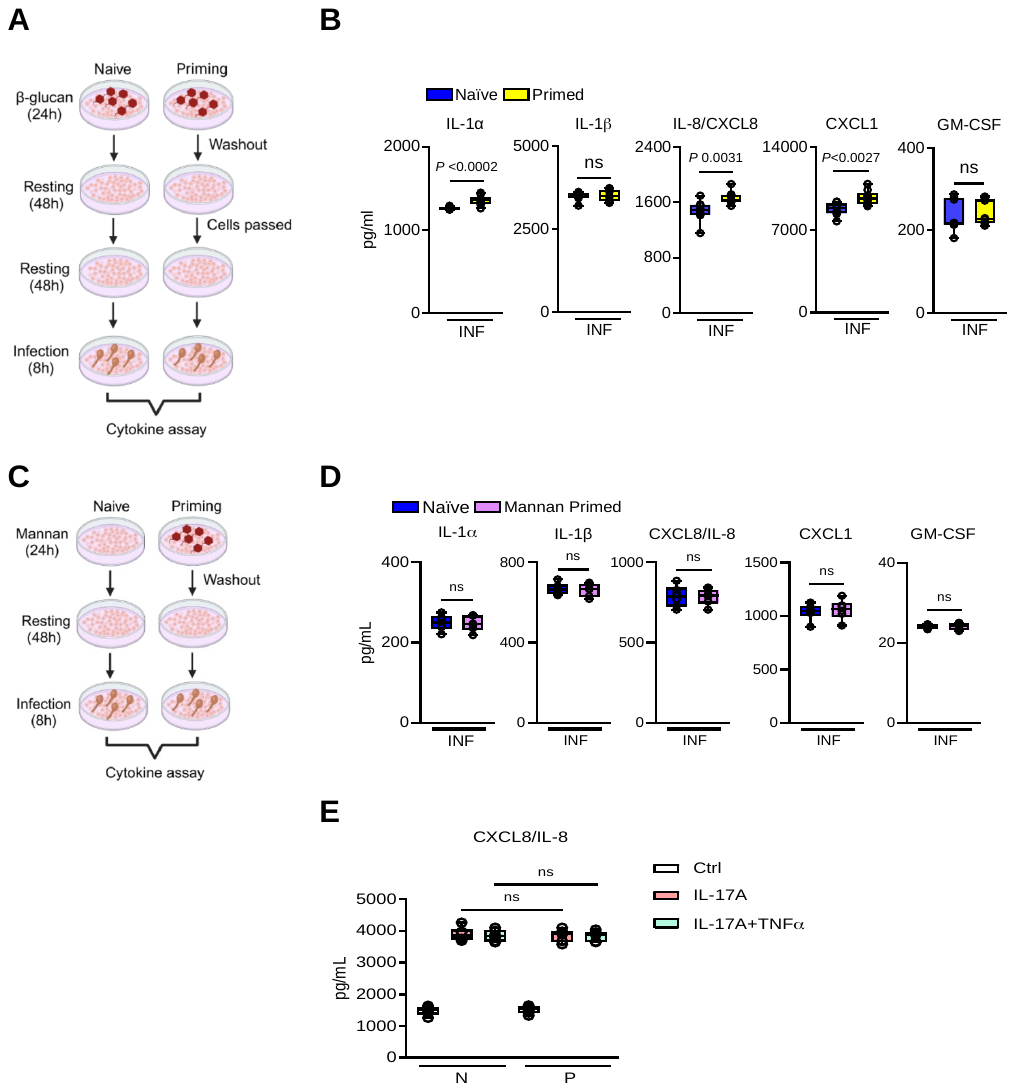

**Figure S5. β-glucan priming enhances epithelial memory after 7 days.** (Related to Figure 2) **A.** Schematic of the *in vitro* experimental design for the induction of long-term β-glucan training. Created with BioRender.com **B.** Levels of proinflammatory cytokines in the culture supernatants of naïve and β-glucan primed epithelial cells after 7 days, 8h after infection with *C. albicans. N=6*; Unpaired Student’s t-test**. C.** Schematic representation of the *in vitro* experimental setup for mannan training. Created with BioRender.com **D.** Cytokine levels in culture supernatants of naïve and mannan-primed epithelial cells 8h after infection with *C. albicans. N=6*; Unpaired student’s t-test. **E.** CXCL8/IL-8 levels in culture supernatants of naïve and β-primed OECs after restimulation with IL-17A and IL17A+TNFα for 8h. *N=6*; One-way ANOVA with Tukey's multiple comparisons test. N – Naïve, P – Primed.

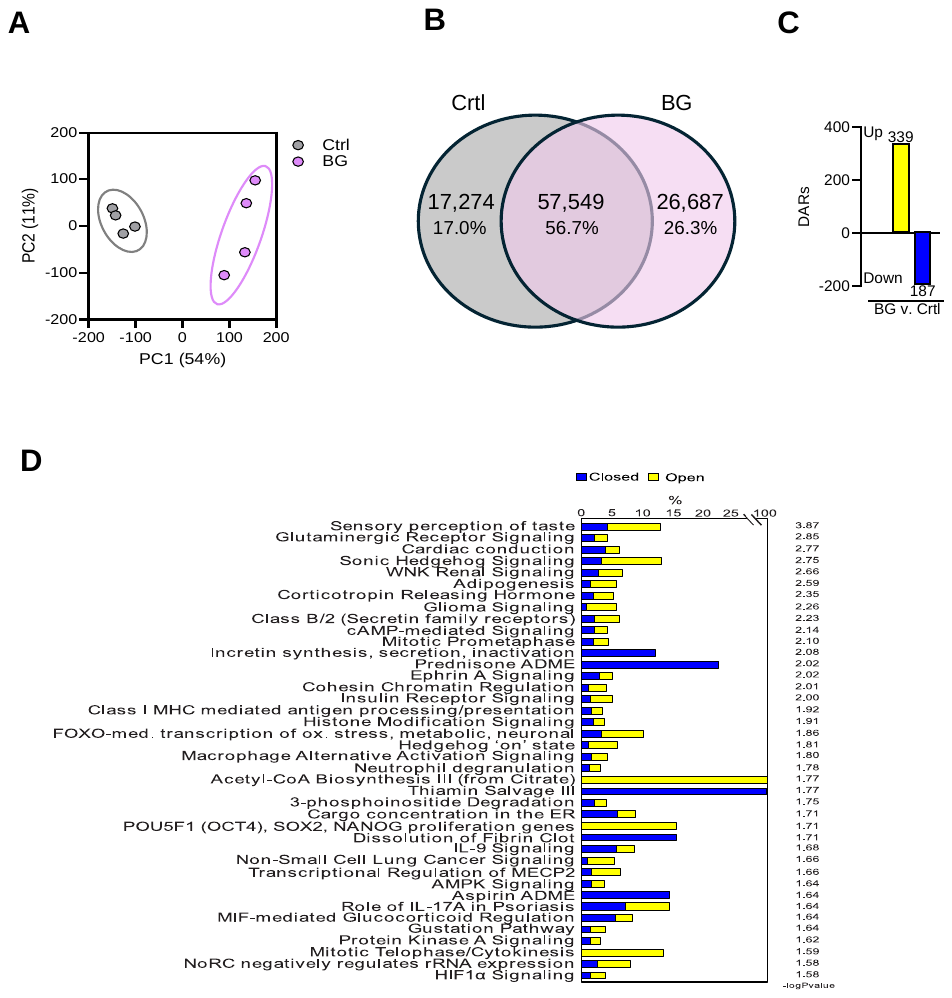

**Figure S6.** **β-glucans recognition alters chromatin accessibility and activates epigenetic and metabolic pathways in oral epithelial cells.** (Related to Figure 2). **A**. Principal component analysis (PCA) of control and β-glucan stimulated epithelial cells for 24h, based on the top differentially accessible loci. Prediction ellipses represent 95% confidence intervals. Each symbol corresponds to an individually sorted subset (*N=4*). **B.** Venn diagram showing overlap of accessible loci between control and β-glucan conditions. **C**. Differentially accessible regions (DARs; p < 0.05) identified by DESeq2, with increased (blue) or decreased (yellow) accessibility depicted by histograms. **D.** Enrichment of canonical pathways in β-glucan-stimulated versus control epithelial cells by Ingenuity Pathway Analysis (n = 4). *N=4*.

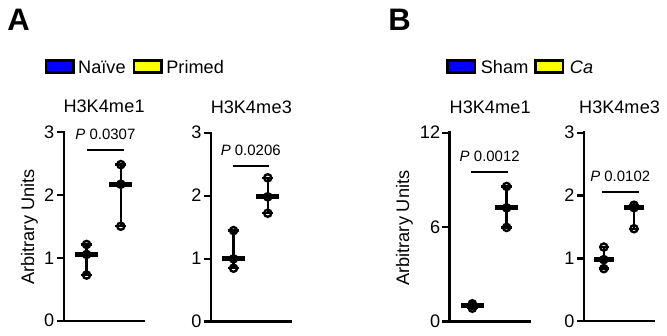

**Figure S7. Quantification of histone methylation changes in epithelial cells and epithelial-enriched tissue. (Related to Figure 2)**. **A.** Densitometric quantification of H3K4me1 and H3K4me3 levels in naïve and β-glucan-primed epithelial cells. *N=3*; Two-tailed Mann–Whitney test. **B.** Densitometric analysis of H3K4me1 and H3K4me3 levels in sham and C. albicans-infected oral mucosal tissue from wild-type mice 5 days post-infection. *N=3*; Two-tailed Mann–Whitney test.

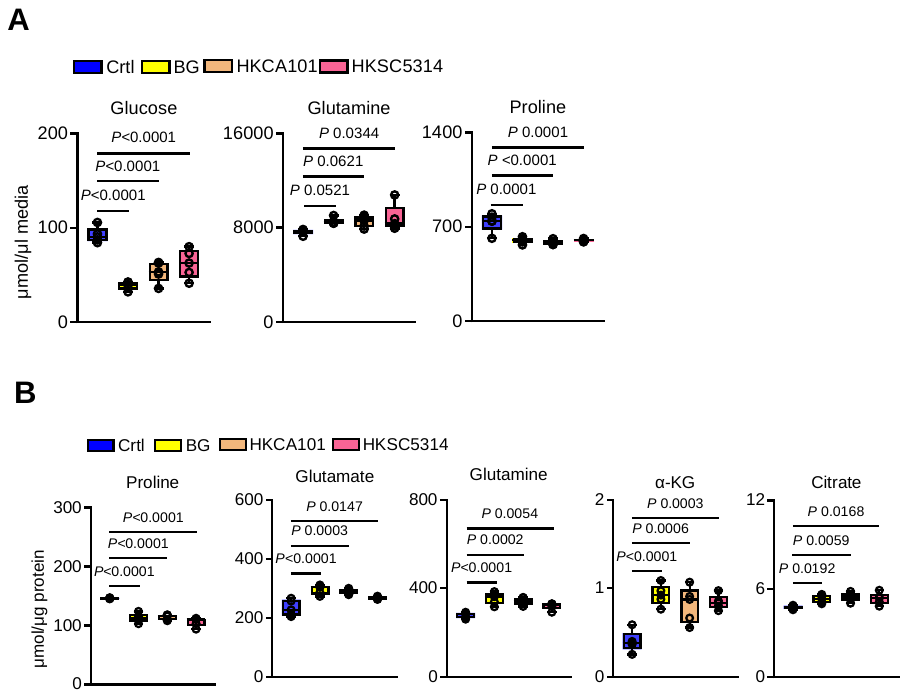

**Figure S8. Exposure to heat-killed *Candida albicans* promotes proline catabolism in epithelial cells.** (Related to Figure 3). **A.** Extracellular glucose, glutamine and proline levels in epithelial cells stimulated with β-glucan or heat-killed *C. albicans* for 24h. *N=5*; One-way ANOVA with Dunnett's multiple comparisons test. **B**. Intracellular metabolite levels in epithelial cells after 24h of stimulation with β-glucan or heat-killed *C. albicans*. *N=5.* One-way ANOVA with Dunnett's multiple comparisons test.

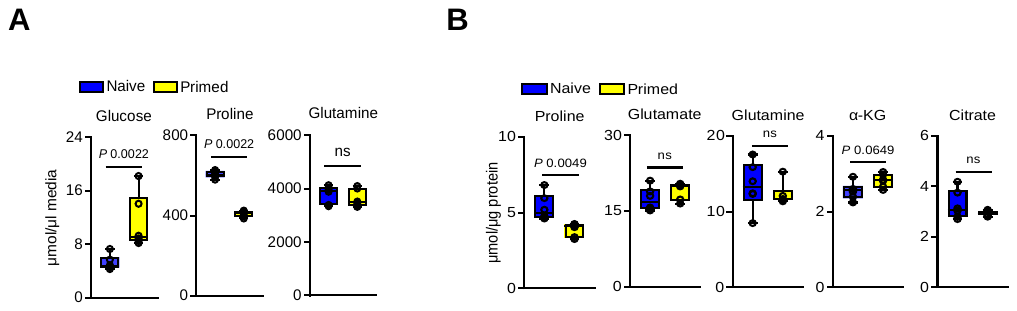

**Figure S9. β-glucan priming promotes proline catabolism in epithelial cells.** (Related to Figure 3).  **A.** Extracellular glucose, proline, and glutamine levels in BG-primed epithelial cells after 48h of rest. *N=6*; Two-tailed Mann–Whitney Test. **B.** Intracellular metabolite levels in BG-primed epithelial cells after 48h of rest. *N=6*; Two-tailed Mann–Whitney Test.

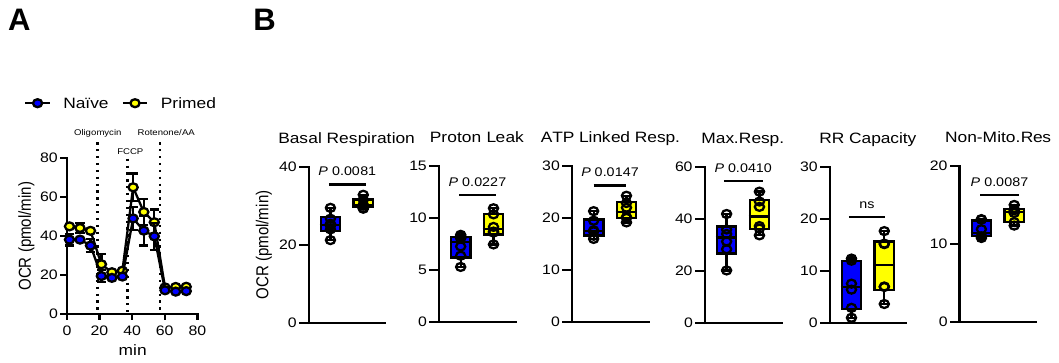

**Figure S10**.**β-glucan priming induces mitochondrial oxidative phosphorylation in epithelial cells.** (Related to Figure 3).  **A.** Seahorse tracings of intact cell respiration in epithelial cells. **B.** Mitochondrial function in naïve and β-glucan-primed epithelial cells after 48h of rest. *N=6*; Unpaired t-test. Basal respiration, proton leak, ATP-linked respiration, maximal respiration, reserve respiratory capacity, and non-mitochondrial respiration were measured.

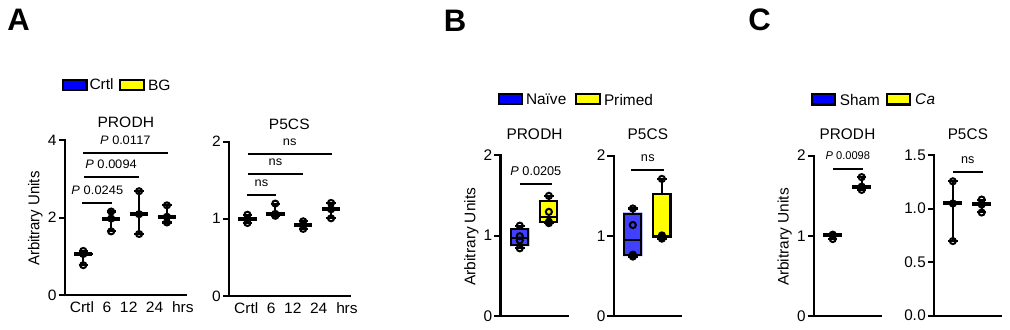

**Figure S11. Quantification of enzymes involved in proline biosynthesis and catabolism in epithelial cells. (Related to Figure 4). A**. Densitometric analysis of **proline dehydrogenase (**PRODH) and **pyrroline-5-carboxylate** synthase (P5CS) expression, normalized to β-actin levels, in control and β-glucan-stimulated epithelial cells. *N=3*; One-way ANOVA with Dunnett’s multiple comparisons test. **B**. Densitometric analysis of PRODH and P5CS expression, normalized to β-actin levels, in naïve and β-glucan-primed OECs. *N=3*; Two-tailed Mann–Whitney test. **C**. Densitometric analysis of PRODH and P5CS expression, normalized to β-actin levels, in sham- and *C. albicans*-exposed oral mucosal tissue from wild-type mice 5 days post-infection. *N=3*; Two-tailed Mann–Whitney test.

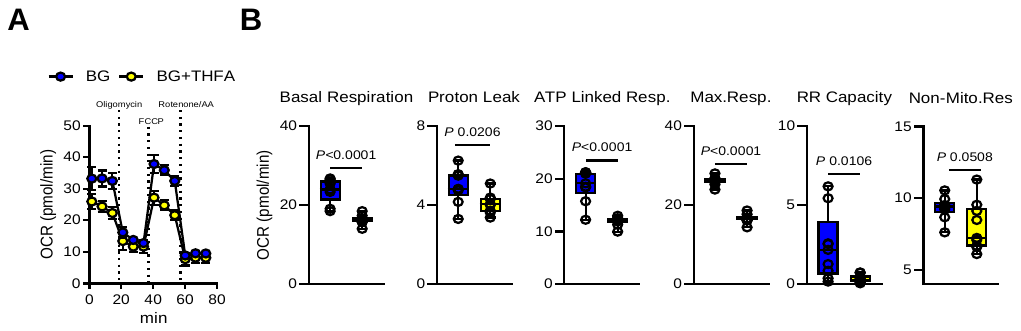

**Figure S12.** **Prodh inhibition impairs mitochondrial oxidative phosphorylation in epithelial cells.** (Related to Figure 4)**. A.** Seahorse tracings of intact cell respiration in human OECs. **B.** Mitochondrial function in β-glucan and THFA exposed to epithelial cells after 24h. *N=6*; Un-paired t-test. Basal respiration, proton leak, ATP-linked respiration, maximal respiration, reserve respiratory capacity, and non-mitochondrial respiration were measured.

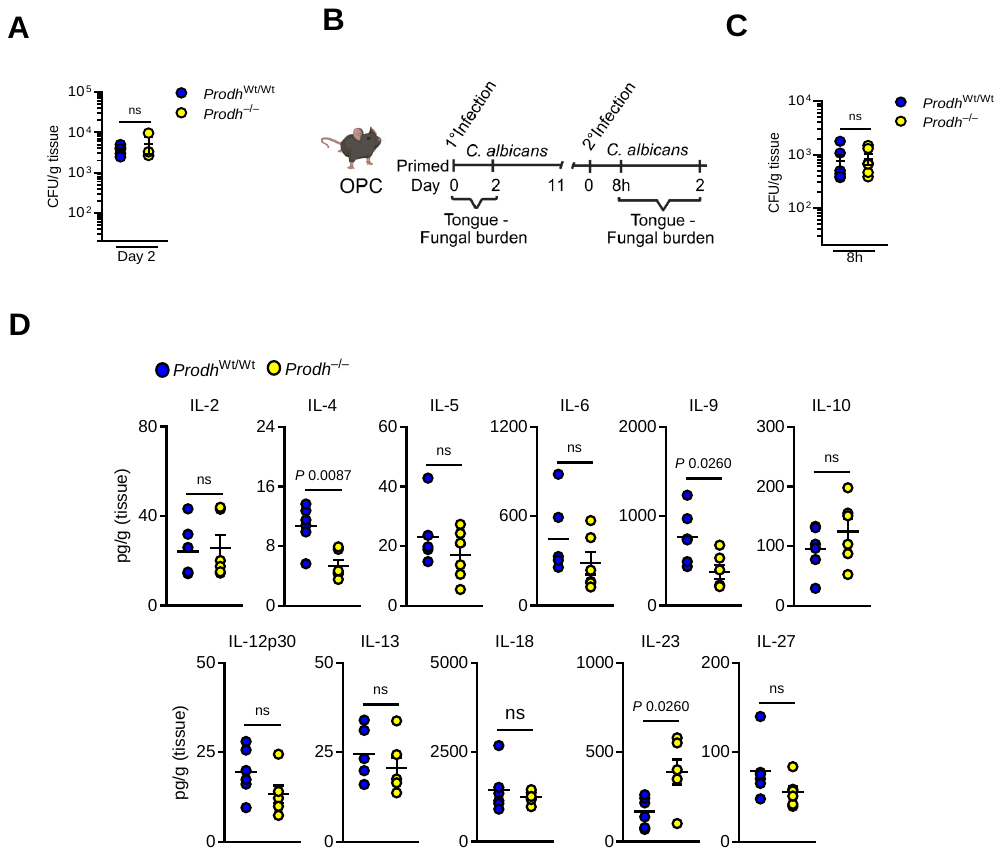

**Figure S13.** **Prodh-deficient mice show reduced distinct inflammatory response during reinfection.** (Related to Figure 4)**. A.** Oral fungal burden in *Prodh*^wt/wt^ and *Prodh^–/–^* mice on day 2 post-primary infection. *N=6*; Two-tailed Mann–Whitney Test. **B.** Schematic of the reinfection model and fungal burden timeline. Created with BioRender.com **C.** Oral fungal burden in *Prodh*^wt/wt^ and *Prodh^–/–^* mice after 8h of reinfection. *N=6*; Two-tailed Mann–Whitney Test. The y-axis represents the limit of detection (20 CFUs/ g of tissue). **D.** Proinflammatory cytokine response in tongue homogenates in wild-type and *Prodh^–/–^* mice after 8h reinfection.

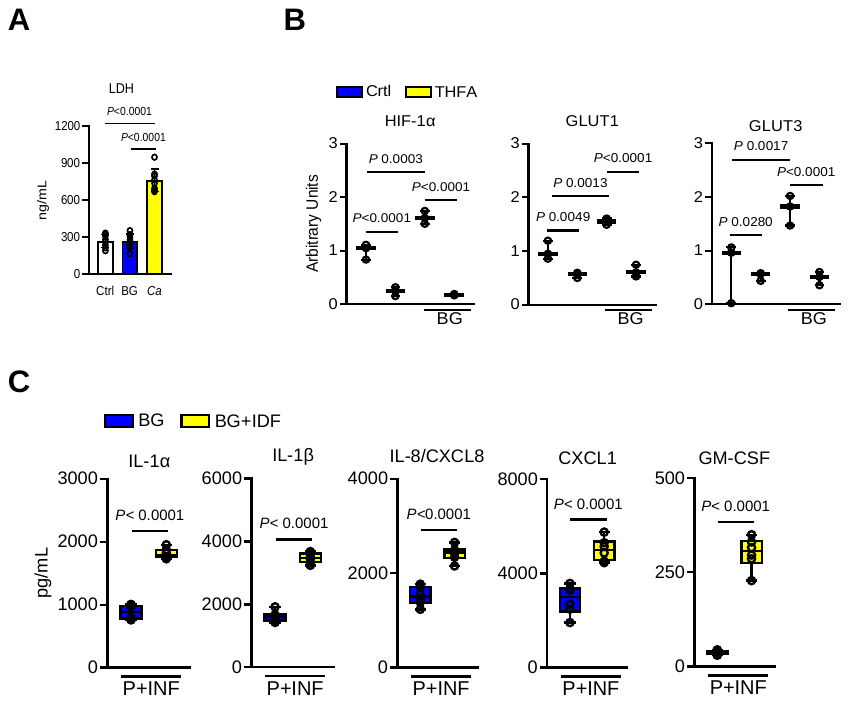

**Figure S14. Lactate dehydrogenase levels remain during β-glucan recognition, and Inhibition of HIF-1α promotes epithelial memory.** (Related to Figure 5)**. A.** Quantification of lactate dehydrogenase release in epithelial cells. **B. Quantification of HIF-1α, Glut1, and Glut3 expression in epithelial cells.** Densitometric analysis of HIF-1α, Glut1, and Glut3 expression normalized to β-actin levels in control, β-glucan and THFA-treated OECs*. N=3*; One-way ANOVA with Tukey's multiple comparisons test. **C.** Levels of chemokines and cytokines in culture supernatants of naïve and β-glucan primed epithelial cells, followed by 8h after infection with *C. albicans. N=6*; Unpaired student’s t-test.

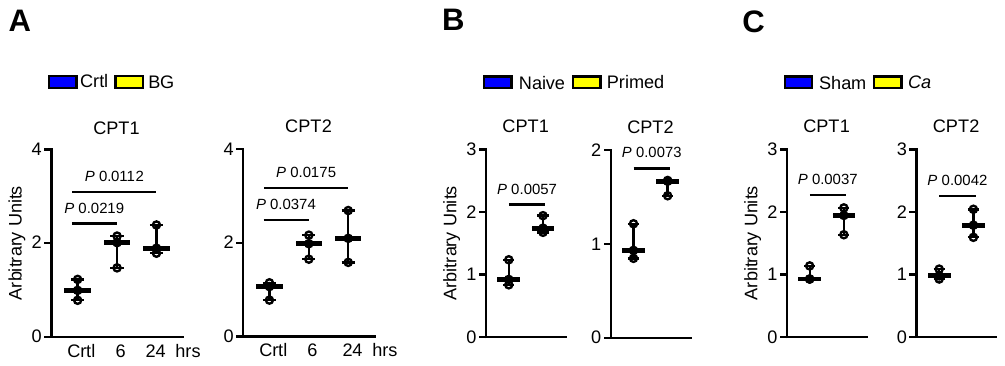

**Figure S15. Quantification of enzymes involved in fatty acid oxidation in oral epithelial cells. (Related to Figure 6)**. **A.** Densitometric analysis of carnitine palmitoyltransferase, CPT1 and CPT2 expression, normalized to β-actin levels, in control and β-glucan-stimulated epithelial cells. *N=3*; One-way ANOVA with Dunnett’s multiple comparisons test. **B.** Densitometric analysis of CPT1 and CPT2 expression, normalized to β-actin levels, in naïve and β-glucan primed epithelial cells after 48h*. N=3*; Unpaired student’s t-test. **C**. Densitometric analysis of CPT1 and CPT2 expression, normalized to β-actin levels, in sham- and *C. albicans*-exposed oral mucosal tissue from wild-type mice 5 days post-infection. *N=3*; Two-tailed Mann–Whitney test.

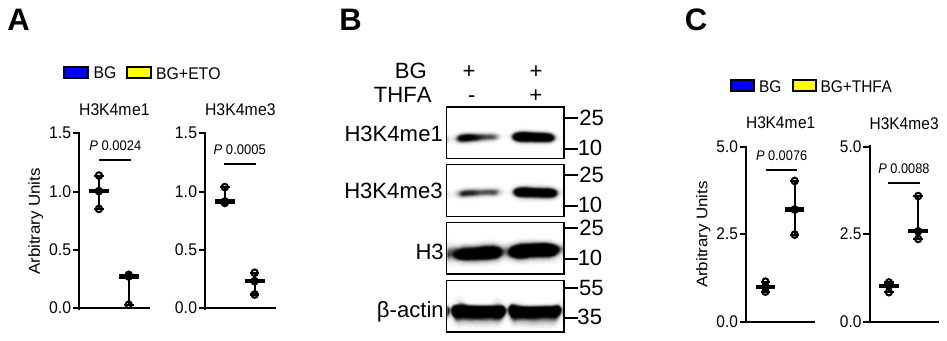

**Figure S16. Quantification of histone methylation in epithelial cells. (Related to Figure 7). A.** Densitometric analysis (arbitrary units) of H3K4me1 and H3K4me3 levels, normalized to histone H3 protein levels, in β-glucan and ETO primed epithelial cells. *N=3*; Two-tailed Mann–Whitney test. **B.** Representative immunoblot showing histone methylation in without or with THFA/BG primed epithelial cells after 48h of rest. C. Densitometric analysis of H3K4me1 and H3K4me3 levels, normalized to histone H3 protein levels, in β-glucan and THFA primed epithelial cells.

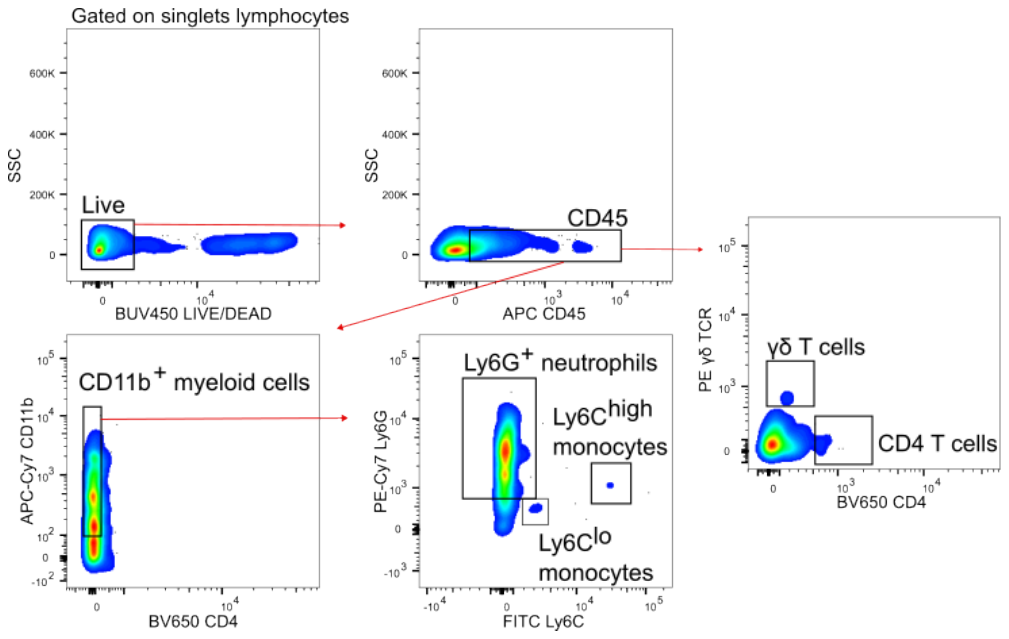

**Figure S17. Gating strategy for immune cell populations in the oral mucosa during reinfection with *Candida albicans*.** Single cells were identified as singlet’s CD45, CD11b^+^, Ly6G^+^, Ly6C^high^, Ly6C^low^, and γδT cells.
